# AI driven approaches in Nanobody Epitope Prediction: Are We There Yet?

**DOI:** 10.1101/2024.10.07.616899

**Authors:** Floriane Eshak, Anne Goupil-Lamy

**Affiliations:** SPPIN CNRS UMR 8003, Université Paris Cité, Paris, France; Biovia Science Council, Dassault Système, Vélizy-Villacoublay, France

**Keywords:** Nanobodies, epitope identification, Benchmarking, structure prediction, deep learning, AlphaFold3, AlphaFold-Multimer

## Abstract

Nanobodies have emerged as a versatile class of biologics with promising therapeutic applications, driving the need for robust tools to predict their epitopes, a critical step for in silico affinity maturation and epitope-targeted design. While molecular docking has long been employed for epitope identification, it requires substantial expertise. With the advent of AI driven tools, epitope identification has become more accessible to a broader community increasing the risk of models’ misinterpretation. In this study, we critically evaluate the nanobody epitope prediction performance of two leading models: AlphaFold3 and AlphaFold2-Multimer (v.2.3.2), highlighting their strengths and limitations. Our analysis revealed that the overall success rate remains below 50% for both tools, with AlphaFold3 achieving a modest overall improvement. Interestingly, a significant improvement in AlphaFold3’s performance was observed within a specific nanobody class. To address this discrepancy, we explored factors influencing epitope identification, demonstrating that accuracy heavily depends on CDR3 characteristics, such as its 3D spatial conformation and length, which drive binding interactions with the antigen. Additionally, we assessed the robustness of AlphaFold3’s confidence metrics, highlighting their potential for broader applications. Finally, we evaluated different strategies aimed at improving prediction success rate. This study can be extended to assess the accuracy of emerging deep learning models adopting a similar approach to AlphaFold3.

## INTRODUCTION

Nanobodies are a unique class of antigen-binding fragments derived from heavy chain only antibodies (HcAbs).^1^ Since their serendipitous discovery in the late 1980s, nanobodies have become versatile tools across a wide range of applications. Understanding the principles governing their selective binding is crucial for advancing rational design approaches, such as affinity maturation and epitope-based nanobody engineering. Deciphering the binding mechanisms of nanobodies, has long relied on structure-based computational tools like molecular docking and homology modeling. Tools like ZDOCK^2^ have shown some success in identifying nanobody epitopes, however, they are highly dependent on the availability and quality of structural templates and require a substantial expertise in molecular modeling.^3,4^ The advent of AI-driven approaches, such as AlphaFold2 has revolutionized the field.^5^ Recently, AlphaFold3 (AF3) marked a major breakthrough, achieving a reported accuracy exceeding 60% in predicting antigen/antibody complexes, a significant improvement over AlphaFold-Multimer v2.3.2 (AF2-M).^5–7^ This has spurred the development of other deep learning models, such as HelixFold-Multimer, OpenFold3 and Chai-1.^8,9^ In this review, we critically evaluate the performance of AF3 and AF2-M in predicting nanobody epitopes, a crucial step in computationally directed nanobody engineering, offering insights into their strengths and limitations.

HcAbs are secreted by Camelidae species, including Llama’s (*Lama glama*), Alpaca’s (*Vicugna pacos*) and Camels (*Camelus dromedarius* and *Camelus bactrianus*).^10^ The antigen-binding domain of HcAbs is also known as single-domain antibodies (variable heavy domain of heavy chain: VHH) or nanobodies, as depicted in Figure 1. Despite their small molecular weight, nanobodies retain full antigen-binding potential and exhibit strong affinity for their targets. ^11,12^ Even though a nanobody’s paratope is smaller than that of conventional antibodies due to the absence of the light chain, it maintains a comparable binding affinity owing to its unique structural features. Structurally, nanobodies consist of a conserved framework (FR1-4) composed of nine antiparallel β-sheets, as depicted in Figure 1. These β-sheets are linked by loops, among which are three hypervariable loops known as complementarity determining regions or CDRs (CDR1-3). The CDRs are crucial for epitope binding making most of the paratope area, with CDR3 being particularly significant due to its variability in sequence composition, amino acids length and loop conformation.

**Figure 1:**
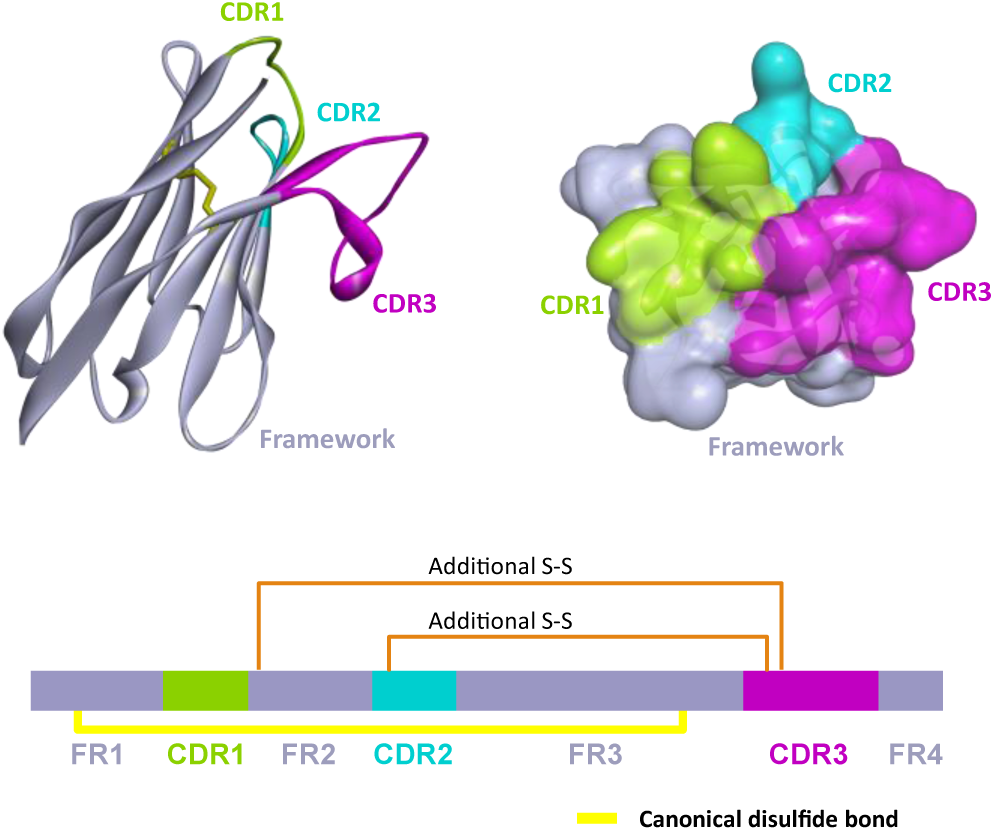
Structural representation of single domain antibodies (VHH or nanobodies). The top left panel displays the nanobody in ribbon representation, with the framework colored in lavender and the three CDRs colored in green, cyan and magenta for CDR1, CDR2 and CDR3, respectively. The top right panel shows a top view of the surface representation of the nanobody, depicting the paratope area, which is mainly formed by the 3 CDRs. The lower panel depicts a sequence illustration, highlighting the distinct regions and the disulfide bridges that connect them: the canonical disulfide bridge between FR1 and FR3 (yellow) and additional disulfide bridges (orange).

The CDR3 structure of VHH is less constrained than its equivalent in conventional antibodies due to the absence of a light chain, allowing it to explore a wider conformational space. These non-canonical loop conformations can be classified based on several criteria, such as the shape of their interaction surfaces into convex, concave or protruded loops.^13^ Recently, a novel classification has been introduced by computing the backbone angles of the C-terminal end into kinked or extended architectures.^14^

To overcome the compact paratope and lack of diversity due to the absence of the light chain, nanobodies have evolved unique features, including an enlarged CDR1 and an extended CDR3, allowing them to target cryptic and hidden epitopes that are inaccessible to conventional antibodies.^15–18^ Additionally, nanobodies feature a conserved canonical disulfide bridge between FR1 and FR3, with some species exhibiting additional disulfide bridges between CDR3 and other CDRs, providing enhanced stability.^10,19^ Overall, nanobodies offer several advantageous properties, such as their compact size, high stability and strong binding affinity, making them ideal for developing next generation biologics.^20–24^

As nanobodies emerge as a promising therapeutic and diagnostic class of biomolecules, understanding their binding modes and interactions with targets can provide valuable insights for affinity maturation and epitope-targeted design.^25–31^ While experimental techniques such as x-ray crystallography, NMR and cryo-electron microscopy can provide this structural information, they are often time consuming and expensive. Therefore, leveraging computational tools that rely on modeling the 3D structures of antibodies and antigens, through homology modeling followed by docking has been used with several successes.^3,4^ However, they are heavily dependent on the availability of suitable templates and require a thorough analysis of the results, making them less accessible to non-expert users.

The development of end-to-end deep-learning based tools has made molecular modeling accessible to non-domain specialists, offering an effective alternative. The most recent AF3 exhibits high performance, with reported highly accurate antigen/antibody complexes predictions exceeding 60% when sampling 1000 seeds.^7,32^ Despite these advances, epitope identification remains challenging due the dynamic nature of proteins and the hypervariability and flexibility of CDR loops, which are critical for antigen recognition. Specifically, with CDR3 reaching considerable lengths and adopting a wide range of conformations.

This study aims to explore the limitations of these emerging tools, guiding readers in effectively analyzing the results. We evaluate the accuracy of the recently released AF3 in predicting nanobody epitopes compared to AF2-M. We present a comparative analysis of 70 epitope identification cases to assess the performance of AF3 and AF2-M, while uncovering the rules that govern nanobody-antigen recognition.

## RESULTS

### I. SELECTION AND DIVERSIFICATION OF THE BENCHMARKING DATASET

A dataset of 70 unique nanobody/antigen complexes was selected from SabDab and the Protein Data Bank.^33,34^ This dataset was not included in the training sets of either AF3 or AF2-M. To ensure the high quality of molecular interactions, critical for this study, the dataset features a median resolution of 2.15 Å, with resolutions ranging from 1.4 Å to 3.6 Å for complexes involving multiple nanobodies. Antigens with several epitopes, co-crystallized with different nanobodies, were assigned the same PDB name with a numerical suffix. To maintain dataset diversity, several criteria were assessed, including the origins of the nanobodies, their targets and their structural variability, as detailed in Supplementary Information Figure 1a.

#### A. DIVERSITY IN NANOBODY ORIGINS AND ANTIGEN TYPES

In the benchmarked dataset, nanobodies were retrieved from diverse Camelidae species, with Lama glama representing over 50% of the dataset, followed by Vicugna pacos at 30.9%, as shown in Supplementary Information Figure 1a. This distribution is consistent with the SabDab nanobody database^33^ where *Lama glama* accounts for 62.7%, followed by *Vicugna pacos* for 24.7%, followed by the other Camelidae species (Supplementary Information Figure 1b). Given that nanobody characteristics can be influenced by their germlines and species of origin, the benchmarked dataset exhibits variability in their properties.^14,19^ To support this, we analyzed the lengths of the three CDRs according to Chothia numbering scheme across different species.^35^ We found that CDR1 is consistently about 7 residues long across all species whereas CDR2 varies, ranging from 4 to 7 residues with a median of 6 residues long in New World Camelids^36,37^ (*Lama glama* and *Vicugna pacos)* and is almost always 6 residues long in Old World Camelids^36,37^(*Camelus dromedarius* and *Camelus bactrianus)*. CDR3, exhibits considerable variability in residue length among most species. Notably, *Camelus bactrianus* features a CDR3 averaging 18 residues with a median length of 19 residues, differing from the wider range of CDR3 lengths found in other species. These findings are detailed in Supplementary Information Figure 1b and Supplementary Information Table T1.

Since we are investigating epitope identification, the dataset includes a diverse range of antigens with varying functions, structures and physiological characteristics. It features transmembrane proteins, such as transporters, ion channels and receptors, as well as soluble proteins like enzymes and interleukins, which are involved in different physiological responses (Supplementary Information Figure 1a). Additionally, the dataset features conformational epitopes and antigens displaying multiple epitopes complexed with different nanobodies.

This diversity highlights the broad range of nanobody origins, characteristics and epitope structural features, thereby minimizing potential bias in the benchmarking dataset.

#### B. DIVERSE YET CONSISTENT CDR3 PROPERTIES OF THE BENCHMARKING DATASET

Given that CDR3 plays a crucial role in epitope binding and constitutes most of the nanobody paratope and considering its high variability, it was essential to ensure diversity in CDR3 properties, including length and conformation.

The Chothia numbering scheme was employed to annotate CDR3 lengths within the dataset, which range from 3 to 24 residues, as depicted in Figure 2A. This broad range reflects the variability within the benchmarking dataset, which, in turn, influences the diversity of CDR3 conformations. Notably, the distribution of CDR3 lengths in the benchmarking dataset is consistent with that observed in the SabDab dataset. The full dataset exhibits a peak at 19 residues, attributed to a high number of nanobodies targeting G-proteins that were excluded from the dataset due to the presence of multiple epitopes interacting with a single nanobody.

**Figure 2:**
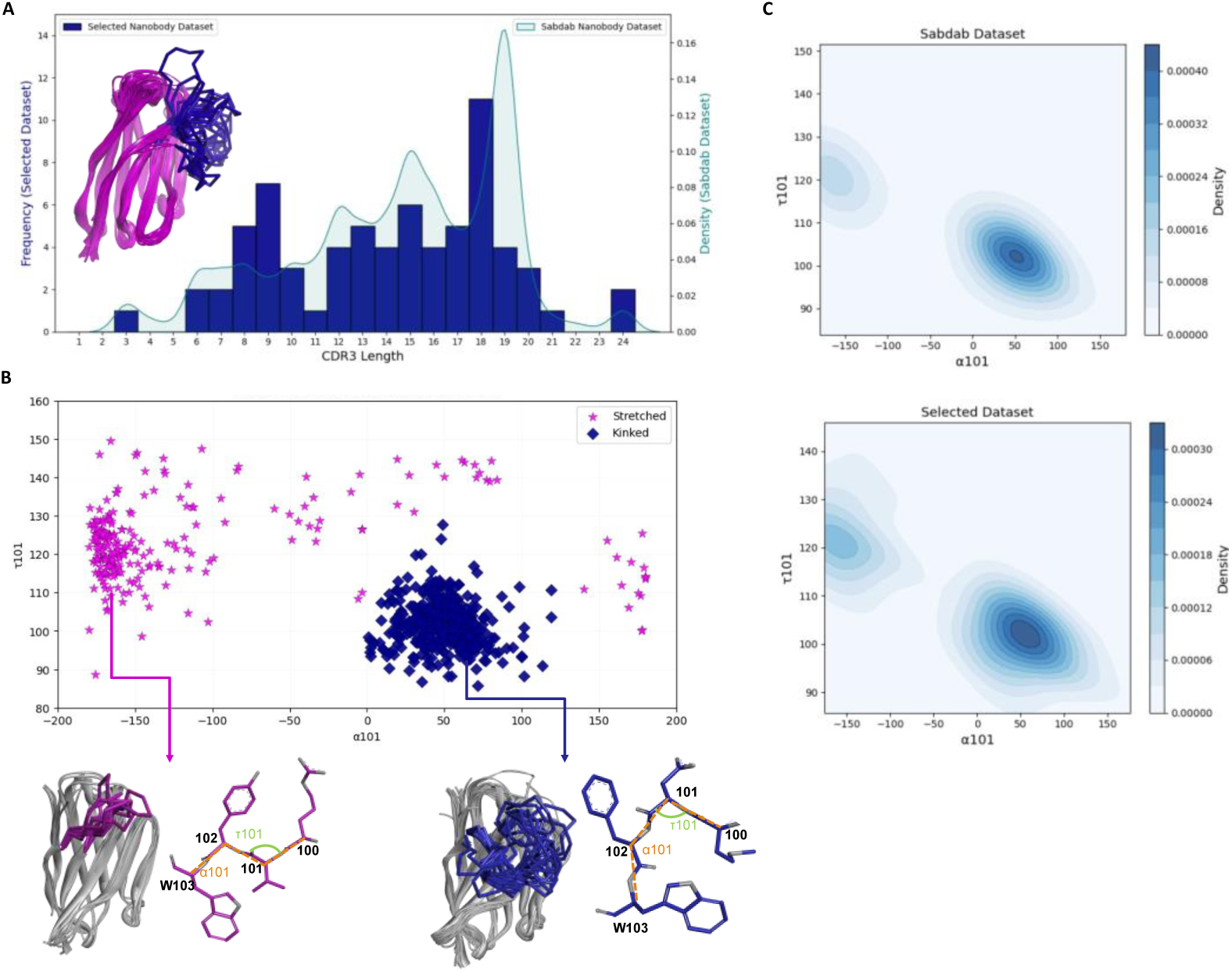
CDR3 characteristics within the benchmarking nanobody dataset. (A) This dual axis plot features a histogram showing the distribution of CDR3 lengths within the benchmarking dataset with a Kernel Density Estimate (KDE) plot representing the CDR3 length distribution in the SabDab nanobody dataset. The x-axis denotes CDR3 length according to the Chothia numbering scheme. The left y-axis represents the number of nanobodies in the selected dataset, while the right y-axis depicts the density of nanobodies in the SabDab dataset. (B) The scatter plot displays the distribution of CDR3 conformations based on their bond and pseudo dihedral angles. The x-axis shows the α101 angle, and the y-axis represents the τ101 angle. The intersection of these angles determines the presence or absence of a kink at the C-terminal end of CDR3 and can provide insights into the conformation of the latter, as either stretched (Magenta) or kinked (Blue). The calculated angles and residues are illustrated for both conformations. (C) Two density plots comparing the distribution patterns of α and τ angles of the SabDab dataset (Upper panel) and the selected dataset (Lower panel). Darker blue areas indicate high density, while lighter blue areas represent lower density. Both plots reveal similar patterns, with a dense, localized area corresponding to the kinked conformation and a less dense cluster representing the stretched conformation

To uncover the impact of CDR3 conformation on shape complementarity and its effect on the binding interaction between a nanobody and its antigen, we adopted the CDR3 classification scheme reported by Bahrami Dizicheh et al.^14^ which distinguishes CDR3 conformations as either kinked or stretched. A kinked conformation is characterized by the CDR3 tip bending towards the nanobody framework, specifically FR2, often forming, a hydrophobic core stabilizing the CDR3 through a triad of hydrophobic contacts.^3^ Conversely, a stretched or extended conformation occurs when the CDR3 loop extends away from the nanobody framework.

To classify these conformations, we followed the method outlined by Bahrami Dizicheh et al.^14^, calculating the pseudo bond and pseudo dihedral angles (τ101 and α101) of the Cα atoms at the C-terminal end of CDR3, where a kink is typically observed.^14,38^ According to the authors and as illustrated in Figure 2B, the angles delineate two distinct CDR3 conformational populations: kinked and stretched.^14^ Notably, the kinked conformation is more localized within certain angle ranges, while the stretched conformation tends to be more dispersed due to its greater flexibility. Consequently, the benchmarking dataset consists of 45 nanobodies with a kinked conformation and 25 with a stretched conformation. To ensure that the benchmarking dataset aligns with the CDR3 conformation distribution pattern observed in the SabDab dataset, we analyzed the density distribution (Figure 2C). Notably, kinked and stretched conformations occupy similar spaces, with kinked conformations displaying a more localized, higher-density cluster, while stretched conformations exhibit a broader range of angle variations.

Based on these findings, we have established that the selected dataset is both diverse and representative of the SabDab nanobody dataset, making it unbiased and suitable for assessing the accuracy of epitope prediction by AF3 and AF2-M.^5,7,33^

### II. EPITOPE IDENTIFICATION ACCURACY

Epitope identification using docking has always been challenging due to its inability to fully account for the structural dynamics of the conformational rearrangement of the paratope and epitope as well as the structural ensembles capturing the dynamics of CDR loops in nanobodies, sampling limitations and its dependence on the quality of structural templates used. AF2-M and later AF3, bypassed these limitations in theory by co-folding the entire complex. These algorithms are capable of predicting protein complexes, including antibody-antigen interactions, but they face certain challenges because antibody-antigen complexes often do not follow the same evolutionary patterns as typical protein-protein interactions. In this study, we will evaluate the performance of both tools in epitope prediction of nanobodies.

#### A. AF3 IMPROVED AF2-M PREDICTIONS REGARDING NANOBODY EPITOPE IDENTIFICATION

The quality of the top-ranked 70 nanobody/antigen complex predictions by AF3 and AF2-M was assessed using DockQ.^39^ DockQ evaluates complex prediction quality by considering the interface root mean square deviation (iRMSD), ligand root mean square of deviation (LRMSD) and fraction of correctly predicted reference contacts (fnat). Based on the predicted and reference models, DockQ classifies predictions into one of four categories: Incorrect, Acceptable, Medium and High quality. DockQ values range from 0 to 1, with higher values indicating better prediction quality, comparable with experimental models (Supplementary Information Table 2).

Across the 70 tested complexes, AF3 achieved correct epitope predictions in 47.1% of cases, with 21.4% considered high-quality predictions, comparable to experimental quality, and 35.7% as intermediate quality. Intermediate quality includes both acceptable and medium quality predictions and indicates that the epitope was successfully identified but with lower accuracy compared to the experimental structure, as illustrated in Figure 3A. This suggests that while AF3 and AF2-M successfully predicted the epitope, they often failed to capture all interactions, exact positioning or the fine-tuned orientation of the nanobody relative to its antigen, as illustrated in Figure 3B. Notably, the high-quality predicted complexes capture the maximum number of interactions, with the orientation of the backbone and side chains closely resembling that of the co-crystal structure. This makes them suitable for direct use in further calculations, as illustrated in Supplementary Information Figure 2. In contrast, intermediate-quality structures require additional modeling techniques to accurately describe the interactions. For medium-quality structures, molecular dynamics simulations can be applied to improve the prediction of interactions. Acceptable-quality predictions, demand more advanced modeling techniques to understand the key interactions between the antibody and antigen. AF2-M correctly predicted nanobody epitopes in 32.8% of cases, with 11.4% considered high-quality and 21.4% as intermediate quality. Consequently, AF2-M failed to predict nanobody epitopes in 67.1% of the cases compared to 52.9% for AF3 (Supplementary Information Table 3).

**Figure 3:**
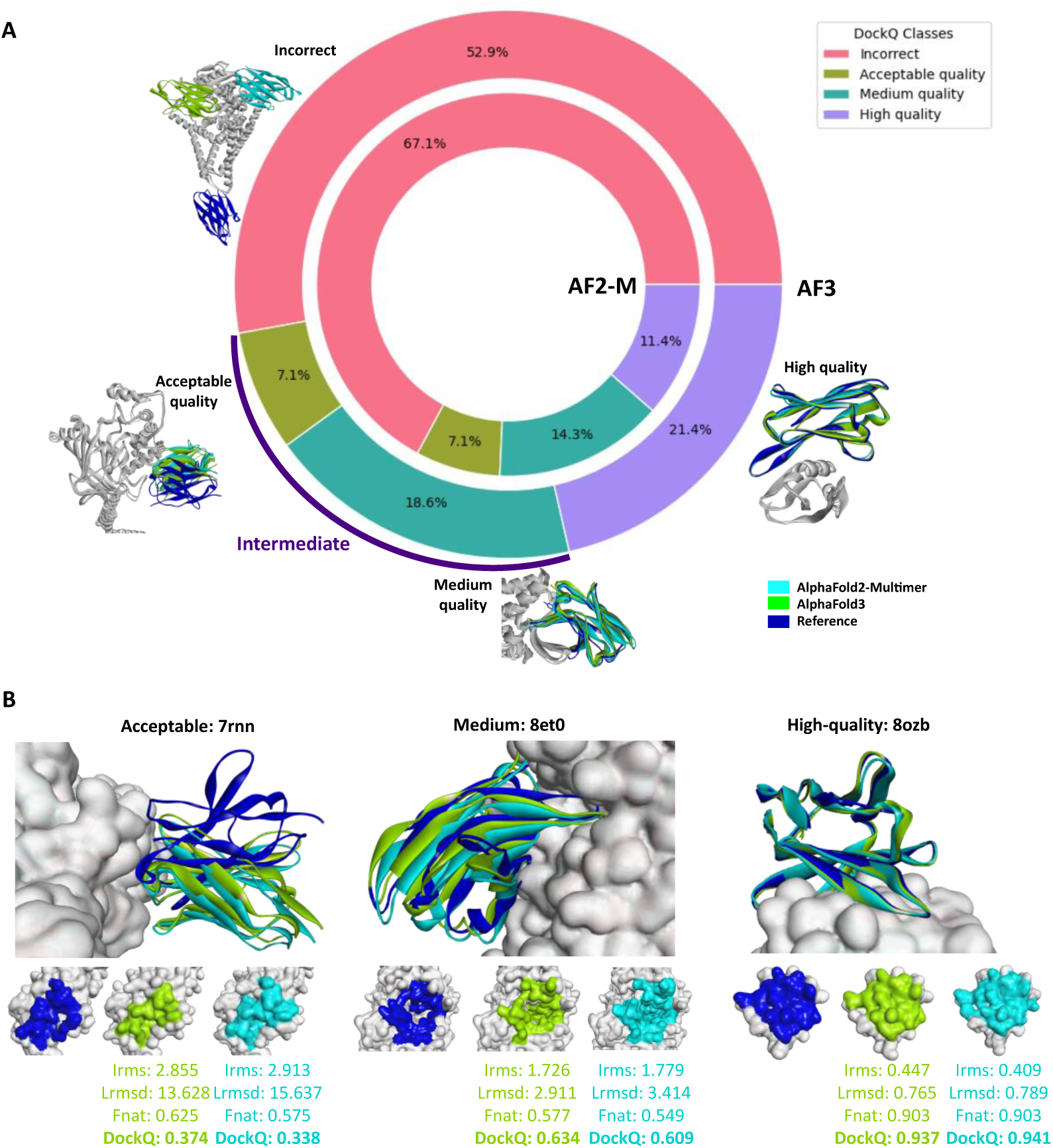
Performance overview of AF3 and AF2-M in nanobody epitope identification. (A) The figure features a dual pie chart, with the outer pie chart representing AF3 and the inner pie chart representing AF2-M. Each chart displays the distribution of DockQ quality classes, indicating the percentage of predictions within each class. Illustrative examples of predicted epitopes belonging to the four distinct DockQ classes are provided, comparing the reference epitope with the epitopes predicted by AF3 and AF2-M. (B) Illustrative examples of correctly identified epitopes across three different DockQ classes: acceptable, medium and high-quality predictions are provided. For each example, the superimposition of the complexes, the epitope imprints and the detailed DockQ score values are shown: irms (interface root mean square of deviation), Lrmsd (ligand root mean square of deviation) and fnat (fraction of correctly predicted reference contacts)

These findings highlight that AF3 has enhanced AF2-M’s performance in epitope identification, increasing the success rate from 32.8% with AF2-M to 47.1% with AF3. Despite this advancement, nanobody epitope identification remains a considerable challenge, as the overall failed predictions still exceed 50%.

### B. VARIABILITY IN PREDICTION QUALITY OF NANOBODY-ANTIGEN COMPLEXES

The DockQ scores revealed significant variability in the prediction quality of both tools. For AF3, DockQ scores ranged from 0.005 in complexes like 8emz^40^, where the predictions failed to identify the epitope, to 0.941 for 8pe1^41^, as depicted in Supplementary Information Figure 3. Similarly, for AF2-M’s DockQ scores ranged from 0.004 for 8sne_1 to 0.941 for 8ozb.^42^ Both algorithms exhibited similar performances in some instances such as in 7ubx, where they both failed to identify the epitope and in 8ozb, where both achieved high quality results. However notable divergences in performance were observed in other instances such as in 8pi_1, where AF3 outperformed AF2-M and in 8q95_1, where AF2-M AF2-M was superior to AF3.

Overall, the significant variability in DockQ scores reflects a non-uniform pattern in epitope structure prediction across the dataset, suggesting that this discrepancy is more likely attributed to various factors.

#### III. KEY FACTORS AFFECTING EPITOPE STRUCTURE PREDICTION ACCURAVY

Given the significant variation in the prediction accuracy of AF3 and AF2-M, we aimed to investigate the factors influencing their performance across the dataset by examining several criteria, including nanobody biophysical properties and spatial representation.

### A. IMPACT OF CDR3 CHARACTERISTICS ON AF3 AND AF2-M PERFORMANCE

Since CDR3 plays a crucial role in nanobody selectivity and antigen recognition, we compared the predictive performance of AF3 and AF2-M by evaluating CDR3’s properties such as its conformation, residue composition and physiochemical characteristics, which can potentially serve as a predeterminant guide for epitope identification quality assessment.

#### 1. CDR3 LOOP LENGTH

Based on the data presented earlier, which illustrates the variability in CDR3 length ranging from 3 to 24 residues, we found that longer CDR3 loops are associated with decreased accuracy in epitope identification for both AF3 and AF2-M, as represented in Figure 4A. This is supported by a notable peak of long CDR3 loops in incorrectly predicted epitopes, with a median CDR3 length of 17 residues. In contrast, a peak shift to the left, indicating a higher density of shorter CDR3 loops, is associated with high-quality predictions, with median CDR3 lengths of 11 residues for AF3 and 12 residues for AF2-M. This suggests that AF3 and AF2-M tend to perform better with nanobodies exhibiting shorter CDR3 lengths. These findings are consistent with the nature of longer CDR3 loops exhibiting greater conformational flexibility, making the prediction of their binding conformation more challenging and thus reducing the accuracy of epitope identification.

**Figure 4:**
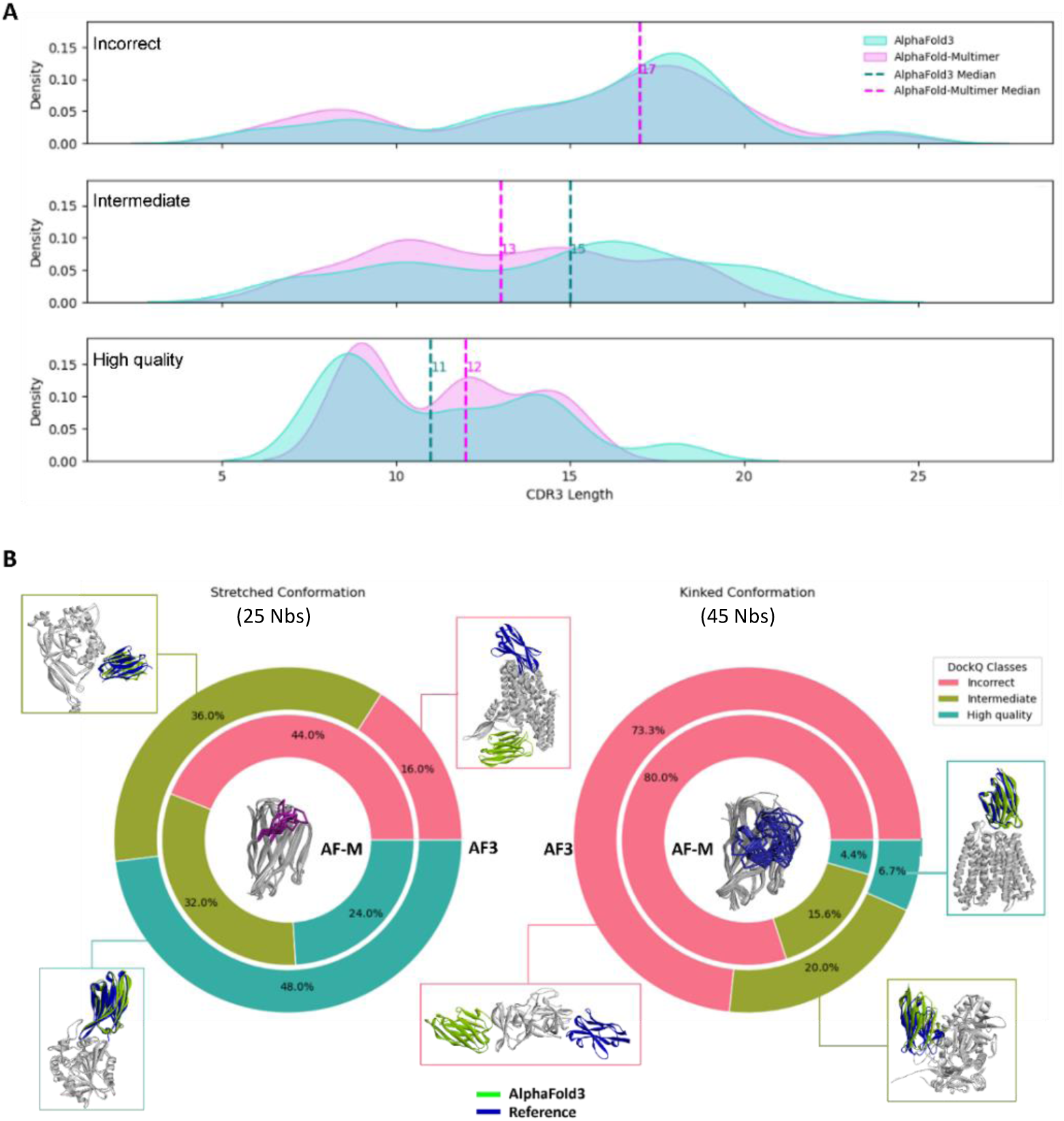
Effect of CDR3 characteristics on epitope identification performances. (A) Effect of CDR3 lengths: The three plots depict the distribution of CDR3 lengths across the three different performance classes for epitope identification. The CDR3 residue lengths are shown on the x-axis and density distribution on the y-axis. AF3 is represented in turquoise and AF2-M in violet. The median CDR3 lengths for each class are also shown. (B) Effect of CDR3 conformation: This diagram shows the percentage distribution of DockQ classes based on the conformation of the CDR3 of the nanobody. The diagram depicts two double pie charts: The outer pie represents the performance of AF3, while the inner pie represents AF2-M. Incorrect poses are colored in pink, intermediate in green and high-quality performances in turquoise. Illustrative examples for each DockQ class are also shown, where the co-crystallized structure is in blue and AF3 is in green.

#### 2. CDR3 LOOP CONFORMATION: KINKED AND STRETCHED

The shape of CDR3 loop is critical in determining the shape complementarity between a nanobody and its antigen, significantly impacting the quality of epitope identification. This is supported by significant chi-square test results (χ² (1) = 22.01, p-value < 0.001, V = 0.55), indicating a strong correlation between the DockQ class and CDR3 conformation class. Our analysis further revealed that both AF3 and AF2-M algorithms perform more accurately when the CDR3 loop adopts a stretched conformation compared to a kinked one, as illustrated in Figure 4B. Specifically, AF3 achieved a correct epitope identification percentage (Intermediate + High quality) exceeding 80% for CDR3’s stretched conformation. In contrast, when the CDR3 loop adopts a kinked conformation, the rate of incorrect predictions exceeded 70%. Notably, AF3 showed a significant improvement over AF2-M in epitope prediction for the stretched conformation, with high quality predictions increasing from 24% to 48%, while the rate of incorrect predictions decreased from 44% to 16%. Despite these improvements in the stretched conformations, kinked conformations remain the majority among existing nanobodies, which significantly impacts the overall success rate of epitope identification.

These findings highlight that improved epitope identification performance is strongly associated with a CDR3 adopting a stretched conformation, which accounts for only 35.7% of the benchmarking dataset. However, since the majority of nanobodies adopt a kinked conformation, accurately predicting their epitopes remains a significant challenge. This emphasizes the crucial role of CDR3 structural conformation in the predictive accuracy of both algorithms and reveals a key limitation of AI-driven approaches in predicting epitopes for nanobodies with kinked conformation.

These findings highlight that improved epitope identification performance is strongly associated with a CDR3 adopting a stretched conformation, which accounts for only 35.7% of the benchmarking dataset. However, since the majority of nanobodies adopt a kinked conformation, accurately predicting their epitopes remains a significant challenge. This emphasizes the crucial role of CDR3 structural conformation in the predictive accuracy of both algorithms and reveals a key limitation of AI-driven approaches in predicting epitopes for nanobodies with kinked conformation.

#### 3. CDR3 RESIDUE COMPOSITION

Upon analyzing the solvent accessible residue composition of the CDR3 and its potential effect on epitope identification performance, we found that for AF3, the percentage of hydrophilic solvent-accessible residues in both stretched and kinked conformations is associated with improved epitope identification predictions, as illustrated in Supplementary Information Figure 4. These findings indicate that a higher composition of hydrophilic residues in the CDR3 loop is linked to better epitope identification performance in AF3. Conversely, a higher proportion of hydrophobic residues is more commonly associated with incorrect epitope predictions. It is important to note that the presence of hydrophobic patch on CDR3 would generally lead to better epitope identification using docking tools such as ZDock.^3^

##### B. IMPACT OF NANOBODY PHYSICO-CHEMICAL CHARACTERISTICS ON AF3 AND AF2-M PERFORMANCE

We analyzed a series of physicochemical properties of the overall structure of nanobodies to assess their potential effects on the epitope identification performance. Specifically, we tested structure-based properties such as the aggregation score, viscosity score, surface charge map scores (QMap), solubility score of the 70 co-crystallized nanobodies. Our findings indicate that there is no correlation between these physiochemical properties and the performance of either AF3 nor AF2-M.

### C. IMPACT OF NATIVE NANOBODY-ANTIGEN BINDING INTERFACE ON AF3 AND AF2-M PERFORMANCE

To assess whether the native binding interface and interactions between the nanobody and its antigen affect the quality of epitope identification, we analyzed several metrics. Our analysis indicates that the surface areas of the antigen and the nanobody may have a slight impact on epitope identification performance, specifically in AF3, as shown in Supplementary Information Figure 5. Our results suggest that larger native surface areas are associated with more accurate predictions by AF3 while no significant effect was observed upon when analyzing the number of native hydrogen bonds or salt bridges between the antigen and the nanobody on the quality of epitope identification.

**Figure 5:**
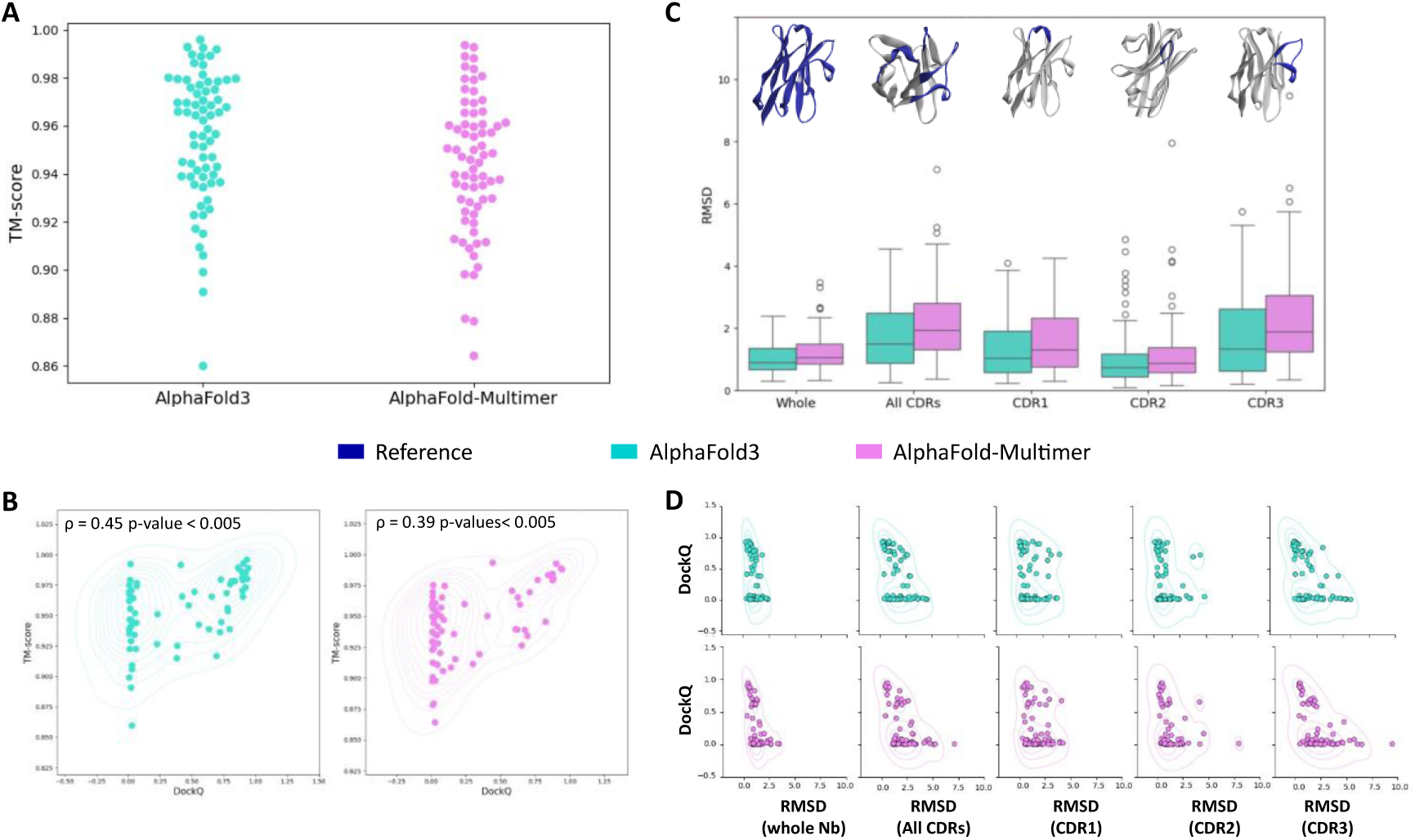
Structural similarity and impact of modeled nanobodies on their epitope identification performances. (A)The swarm plot displays TM-scores (y-axis) for each nanobody (in complex) predicted by AF3 (cyan) and AF2-M (violet). (B) The dual plots represent the distribution of the 70 nanobody (in complex) models, with the y-axis representing the TM-score and x-axis the DockQ score. Overlaid on each scatter plot is a kernel density estimate (KDE) plot indicating areas of higher density. AF3 results are represented in the left panel and AF2-M are shown in the right panel. Notably, high DockQ scores are associated with high TM-scores. However, nanobody models with low DockQ scores exhibit a wide range of TM-scores. Spearman correlation coefficients and their corresponding p-values are shown. (C) The Boxplot displays RMSD values (Å) (y-axis) for different nanobody regions: whole nanobody (CDRs + Framework), combined CDRs, CDR1, CDR2 and CDR3 (x-axis). AF3 data is shown in turquoise and AF2-M data is in violet. Designated nanobody regions are displayed at the top of the plot in blue. (D) The scatter plots show RMSD values (Å) for different nanobody regions on the x-axis versus DockQ scores on the y-axis. The top panel represents AF3 values (in turquoise) and bottom panel represents AF2-M (in violet). Each scatter plot is overlaid with a kernel density estimate (KDE) plot to highlight areas of higher density, with denser regions surrounded by more circles.

### D. EVALUATING THE EFFECT OF NANOBODY STRUCTURE PREDICTIONS ACCURACY ON EPITOPE IDENTIFICATION PERFORMANCE

To evaluate the quality of predicted bound nanobody models, we assessed several factors that define the shape of the paratope and influence its interaction with the antigen. These factors include structural similarity between nanobody models and the relative co-crystal structures, predicted CDR3 loop conformation, the secondary structure of the CDR3 loop and their ability to predict the presence conserved disulfide bridge.

#### 1. STRUCTURAL ACCURACY OF NANOBODY PREDICTION: CRITICAL ROLE OF CDR3 ON EPITOPE IDENTIFICATION PERFORMANCE

While crystal structures represent only one of many possible conformational ensembles of a complex, they provide insights into stable spatial rearrangements formed upon epitope and paratope binding. The structural similarity of the predicted nanobody was assessed using the template modeling score (TM-score)^43^.

As displayed in Figure 5A, both AF3 and AF2-M demonstrated high accuracy in bound nanobody structure prediction, with median TM-scores of 0.963 and 0.946, respectively. To assess whether nanobody model quality affects epitope prediction, we found that despite a weak to moderate Spearman correlation between TM-score and DockQ, high DockQ values are consistently associated with high TM-scores, as illustrated in Figure 5B. This is further supported by the presence of a dense cluster of nanobodies with both high TM-scores and high DockQ values for AF3 and AF2-M. Notably, no instances of low TM-scores paired with high DockQ scores were observed. However, several cases show low DockQ values paired with high TM-scores, suggesting that while high-quality nanobody models are important for accurate epitope prediction, nanobody quality alone does not ensure successful epitope identification Given that TM-scores evaluate the entire nanobody structure, they are influenced by the conserved framework regions, which are often well predicted by computational tools. To assess structural accuracy, the RMSD of the three CDRs was examined, as depicted in Figure 5C. AF3 and AF2-M exhibited comparable performances, with AF3 consistently showing slightly lower RMSD values. Among the CDRs, CDR2, typically composed of 6 residues, exhibited the lowest RMSD medians (0.735 Å for AF3 and 0.89 Å for AF2-M), indicating its high prediction accuracy. In contrast, CDR1, despite being in average 7 residues long, showed higher RMSD medians (1.044 Å for AF3 and 1.306 Å for AF2-M), likely due to its secondary structure variability. As for CDR3, known for its significant diversity in sequence lengths, secondary structures and conformational variability, it exhibited the highest RMSD medians (1.34 Å for AF3 and 1.902 Å for AF2-M).

To further assess the impact of CDRs structure prediction on epitope identification, the correlation between CDR RMSD values and DockQ scores was analyzed. Spearman correlation coefficients revealed that CDR3 loop modeling had a significant impact on DockQ scores for both AF3 and AF2-M (Supplementary Information Table 3). Additionally, lower RMSD values for CDR3 are associated with higher DockQ scores (Figure 5D), suggesting that while an accurate epitope predicted models relied on a high-quality CDR3 model, it does not guarantee success.

In conclusion, while both AF3 and AF-2 demonstrate robust nanobody structure prediction, CDR3 modeling remains challenging but crucial for effective epitope identification. This is consistent with the diversity of CDR3, constituting a significant area of the paratope surface, thereby greatly influencing epitope recognition.

#### 2. CDR3 CLASS PREDICTION (KINKED/STRETCHED) ACCURACY AND ITS EFFECT ON EPITOPE PREDICTION

Given the crucial role of CDR3 architecture in nanobody selectivity and antigen binding, accurately predicting its folding (kinked or stretched) is essential. To evaluate the accuracy of CDR3 folding prediction, we compared the τ101 and α101 angles of the predicted models, which revealed consistent patterns with experimental structures (Supplementary Information Figure 6a and 6b). Both models successfully predicted kinked and stretched conformations, although AF2-M misclassified one kinked conformation into a stretched conformation.

**Figure 6:**
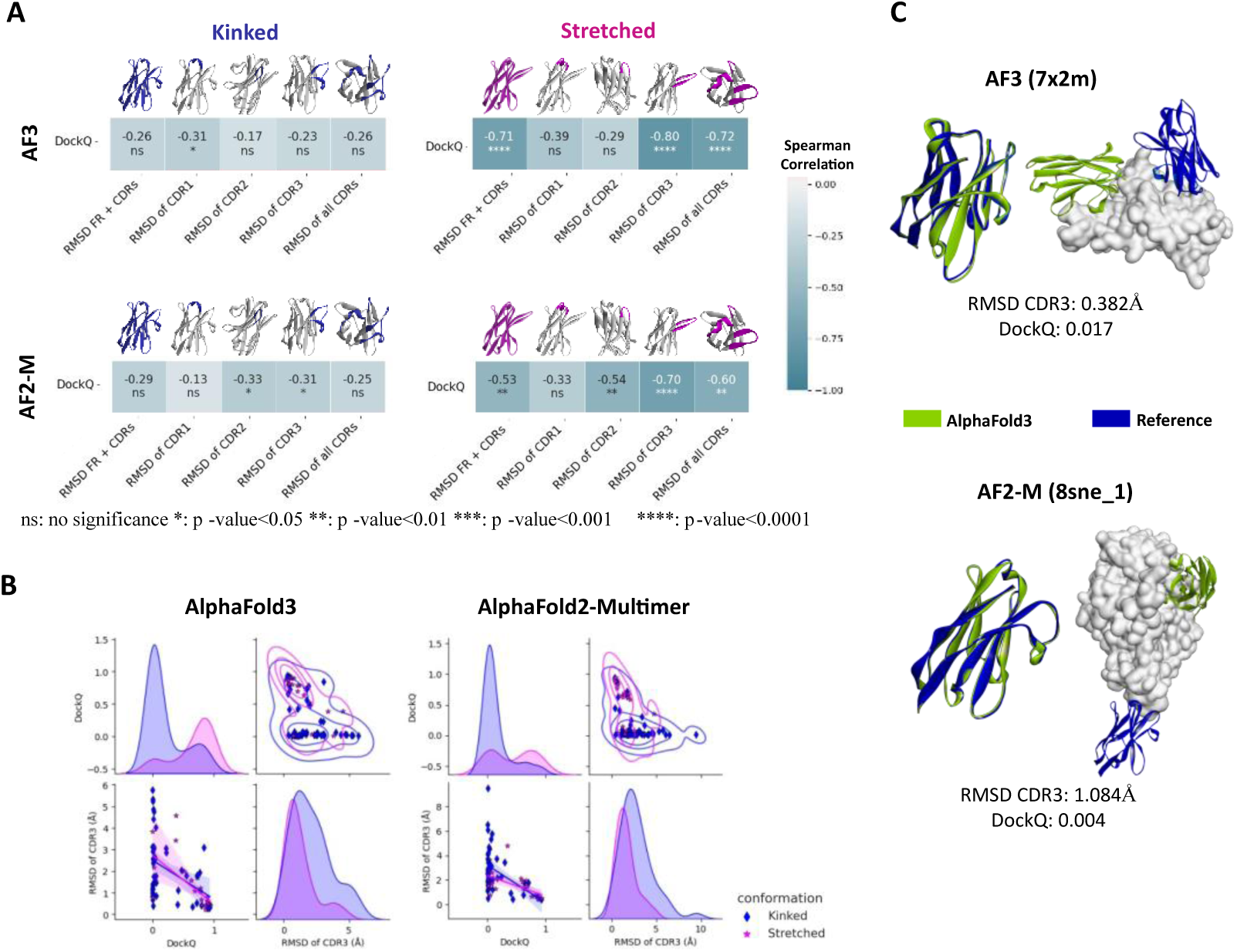
Impact of structural similarity of nanobody models on epitope identification quality across the different CDR3 conformations. (A)The Spearman correlation coefficients between DockQ scores and the structural similarity degree of the different nanobody regions (RMSD). The left panel corresponds to the kinked conformation and the right panel corresponds to the stretched conformation. The Upper panel shows results of AF3 and the lower panel for AF2-M. The color gradient ranges white (no-correlation) to teal (strong anti-correlation). P-values are provided to indicate the statistical significance of the results. (B) Pair plots represent the distribution of RMSD of CDR3 versus DockQ scores for both stretched (magenta) and kinked (Blue) conformations. The left Panel represents AF3 results while the right panel shows AF2-M results. Upper plots include KDE plots to highlight areas of higher density, with denser regions surrounded by more circles. Lower plots include regression lines with bands indicating the 95% confidence for the regression estimates. (C) Illustrative examples of nanobodies adopting the kinked conformation with low DockQ values amid exhibiting low CDR3 RMSD values. The upper panel shows AF3 results while the lower panel displays AF2-M. AF3 structures are colored in green and the reference structures are shown in Blue. There are no instances where CDR3 adopting the stretched conformation was well predicted yet showed failed epitope identification, thus no illustrative example is available.

Since the CDR3 shape determines how nanobodies interact with their antigens, we found that CDR3 model accuracy and epitope identification performance are highly dependent on CDR3 shape. For kinked conformations, the median RMSD of the CDR3 backbone is significantly higher than for stretched conformations (Supplementary Information Table 5). This discrepancy highlights the challenges in predicting kinked CDR3 conformations, reflecting its potential impact on epitope identification quality.

Our analysis revealed that a high-quality CDR3 model in the stretched conformation is strongly associated with accurate epitope identification performance, unlike kinked conformations.

This is supported by a strong Spearman correlation between DockQ scores and RMSD values for stretched CDR3 conformations, while kinked conformations exhibited a weak correlation (Figure 6A). Figure 6B further highlights this distinction, where the densest cluster in kinked conformations corresponds to low RMSD values paired with low DockQ scores. In contrast, for stretched conformations, the densest cluster aligns with low RMSD values and high dockQ scores. Additionally, Figure 6C illustrates instances where, despite accurate CDR3 modeling in the kinked conformation, epitope prediction fails. In contrast, such cases were not observed for the stretched conformation, where accurate CDR3 models consistently resulted in successful epitope identification.

These findings suggest that while successful epitope identification relies on the accuracy of the CDR3 model, a high-quality stretched CDR3 model significantly enhances the likelihood of accurate epitope identification. Conversely, the quality of kinked CDR3 modeling alone does not ensure effective epitope identification.

#### 3. CDR3 SECONDARY STRUCTURE PREDICTION ACCURACY AND ITS IMPACT ON EPITOPE IDENTIFICATION

Due to the significant variability in residue composition, length, and conformation of CDR3 loops, they can exhibit diverse secondary structure compositions. In kinked conformations, longer CDR3 sequences frequently feature one or more 3-10 helix turns (Figure 7A). Conversely, stretched CDR3 loops are often characterized by the presence of two connected β-sheets.^14^ This variation in secondary structure highlights the distinct structural motifs associated with different CDR3 sequences and conformations, which can affect the orientation of side chains and consequently, the surface of the loops which will have a direct impact on both the model then epitope prediction (Figure 7B).

**Figure 7:**
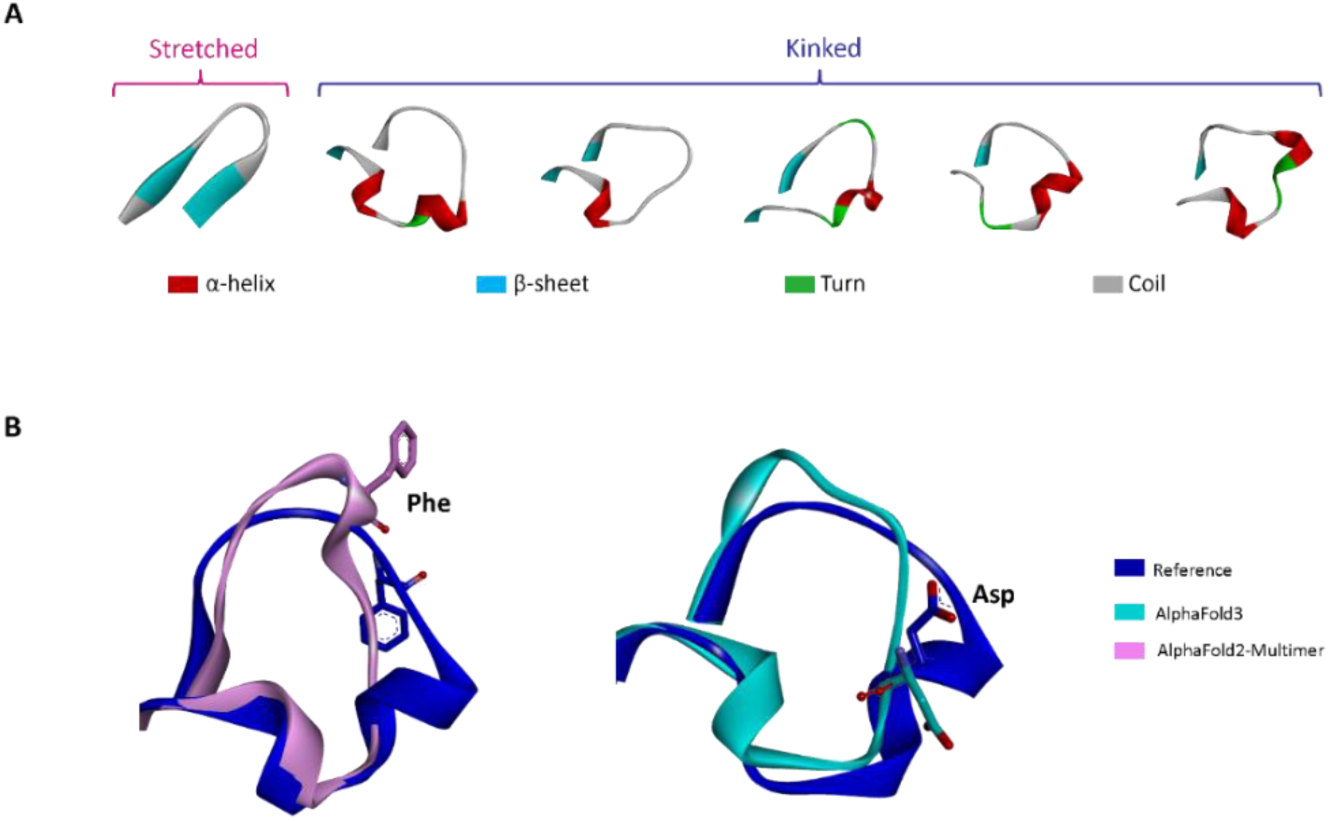
Secondary structure variability within CDR3 loop and its effect on side chains orientation. (A) This figure illustrates the variability in secondary structure composition within CDR3 loops, specifically comparing the stretched and kinked conformations. The kinked conformation shows more secondary structure variability, including the presence of one or more helices positioned at the tip, N-terminal or C-terminal end of CDR3 loop. (B) Two illustrative examples demonstrating how different secondary structures result in varying side chain orientations. The left panel shows the superimposition of the AF2-M model (pink) with the co-crystal structure (blue). The presence of a shifted helix in the N-terminal side of the CR3 loop in AF2-M model leads to a different orientation of the Phe side chain (sticks) when compared to the co-crystal structure. The right panel displays the superimposition of the AF2-M model (cyan) with the co-crystal structure (blue). The absence of a helix at the tip of CDR3 in the AF3 model, compared to the co-crystal structure, results in a different orientation of the Asp side chain.

To further evaluate CDR3 model quality beyond atomic coordinates and predicted conformations, we calculated the percentage of secondary structure retrieval within the CDR3 loops (Supplementary Information Table 6). Notably, AF3 outperformed AF2-M, achieving a median retrieval percentage of 91.99% for overall structures compared to 83.77% for AF2-M (Supplementary Information Figure 7a). This highlights AF3’s ability to provide more accurate secondary structure predictions for CDR3. Notably, AF3 excelled in predicting stretched conformations with a median retrieval percentage of 100% compared to AF2-M’s 87.5%, reflecting its superior performance in modeling stretched conformations. In contrast, the high secondary structure diversity of kinked conformations, renders their structure prediction more challenging.

To assess the impact of accurate secondary structure prediction on epitope identification, we analyzed the secondary structure retrieval percentage across different DockQ classes (Supplementary Information Figure 7b). Both AF3 and AF2-M demonstrated that models with high epitope identification quality exhibited better alignment of secondary structure motifs with the co-crystallized structure. High DockQ scores were consistently associated with a high degree of conserved secondary structure in CDR3 loops, suggesting the importance of a conserved secondary structure for accurate epitope identification.

In conclusion, AF3 improved AF2-M performance in CDR3 modeling by enhancing the accuracy of secondary structure predictions, particularly in stretched conformations. Moreover, in high-quality complex models, the secondary structure is more accurately predicted, which is critical for shape complementarity and strongly correlates with higher prediction accuracy.

#### 4. DISULFIDE BRIDGE PREDICTION AND IMPACT ON EPITOPE IDENTIFICATION

As discussed earlier, nanobodies are highly stable and can withstand a wide range of pH and temperatures. This stability is mainly due to a conserved disulfide bridge between the framework regions FR1 and FR3, located near the N-terminal end of the canonical CDR3. Some nanobodies may exhibit an additional disulfide bridge linking CDR1 or CDR2 to CDR3 (non-canonical), further stabilizing the structure of CDR3.^10,19^

In our analysis, AF2-M outperformed AF3 in predicting canonical disulfide bonds, with a retrieval rate of 80.35% compared to 67.85%, relative to the total number of canonical disulfide bridges present in the co-crystal structures (Supplementary Information Figure 8). Both algorithms exhibited limitations in identifying non-canonical disulfide bridges with AF2-M predicting 2 out of 10, while AF3 failed to predict any. This highlights AF3’s limitations in predicting the accurate orientation of conserved cysteines responsible for both canonical and non-canonical disulfide bridges. Despite the importance of disulfide bridges for nanobody stability, their prediction did not correlate with improved epitope identification performance in either AF3 or AF2-M.

**Figure 8:**
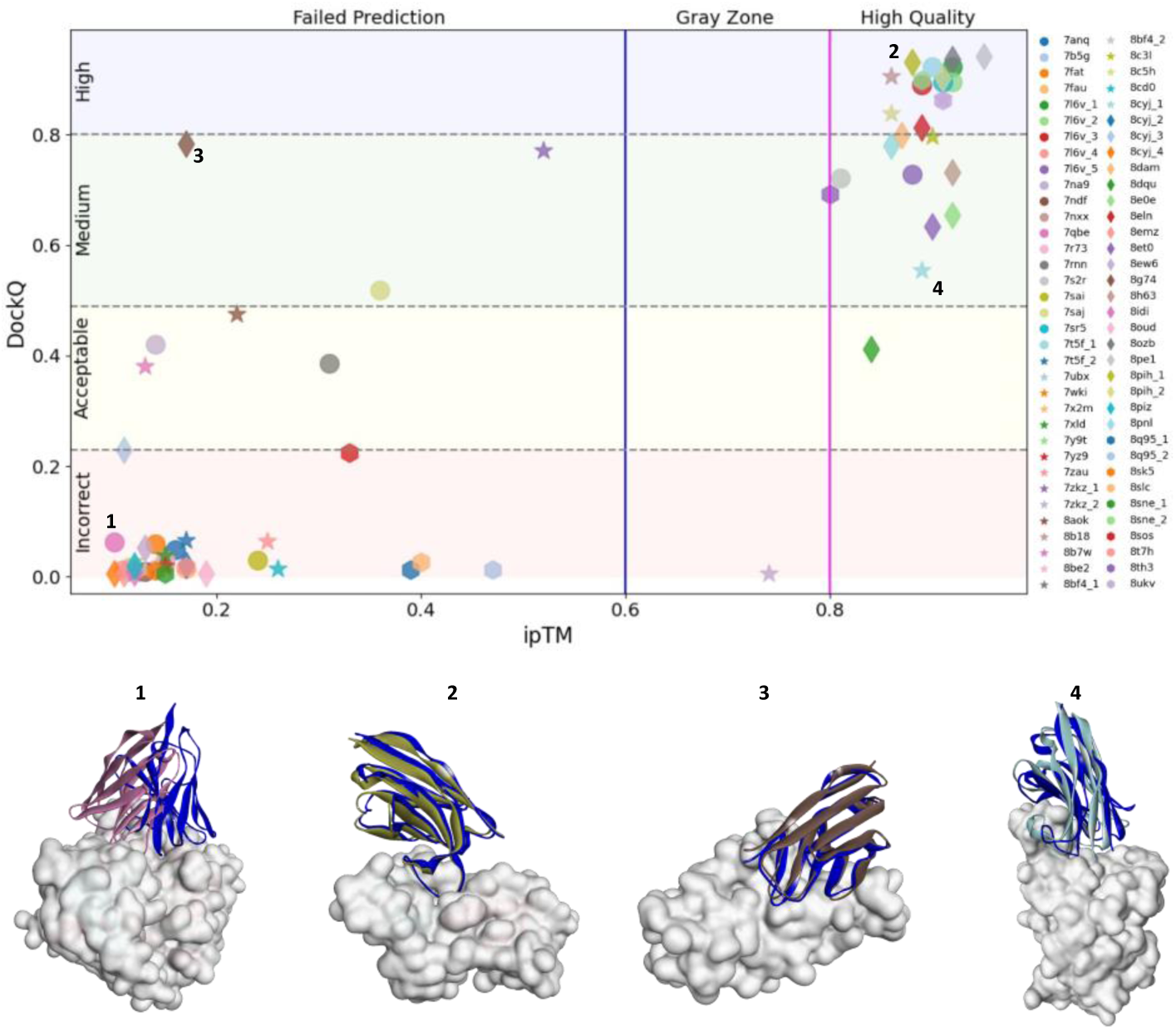
AF3’s ipTM score accuracy in nanobody epitope identification. This figure depicts the correlation between ipTM scores (x-axis) and DockQ scores (y-axis). The upper panel shows each complex example plotted based on its DockQ score and its ipTM score produced by AlpaFold3. DockQ scores are divided into 4 classes: incorrect, acceptable, medium and high quality, while ipTM scores are classified into failed prediction, gray zone and high quality. Each PDB ID is represented using a unique combination of shape and color. Illustrative examples of accurate predictions: true negative (1), true positive (2) and inaccurate ipTM scoring (3,4) are shown.

### IV. ACCURACY OF AF3 CONFIDENCE SCORE AND ITS POTENTIAL APPLICATION

Similar to docking tools, which use scoring functions to rank poses and ideally indicate the likelihood of being close to the native structure, AF3 and AF2-M generate confidence scores that reflect the accuracy and reliability of the predicted structure, while also serving to rank models. AF3’s confidence scores include pTM (predicted template modeling) which measures the accuracy of the predicted structure of a complex by estimating the TM-score for a superposition between the predicted model and the hypothetical true structure and its variant, the ipTM (interface predicted template modeling), which evaluates the accuracy of the predicted interface in a protein-protein complex.^5^ Both ipTM and pTM are dependent on the predicted alignment error (PAE) matrix which reflects the degree of certainty of a model regarding the relative position of residue pairs within the predicted structure. In this study, we will explore the accuracy of these confidence scores and their potential applications.

### A. EVALUATING IPTM AS SELF-ASSESSMENT METRIC FOR AF3 EPITOPE PREDICTION

Given that the goal of this study is to assess the epitope identification accuracy of AF3, it is crucial to evaluate the reliability of ipTM as a self-assessment metric. An ipTM score below 0.6 indicates a failed prediction, while a score above 0.8 suggests a high-quality prediction. Scores between 0.6 and 0.8 indicate uncertainty in the prediction.^5^ As shown in Figure 8, ipTM accurately assessed AF3’s prediction quality in 70% of cases. Additionally, a strong Pearson correlation coefficient of 0.88 (p-value<0.001) was observed between the DockQ and ipTM values, reinforcing the reliability of AF3 in identifying near-native nanobody epitopes (Supporting Information Figure 9).

**Figure 9:**
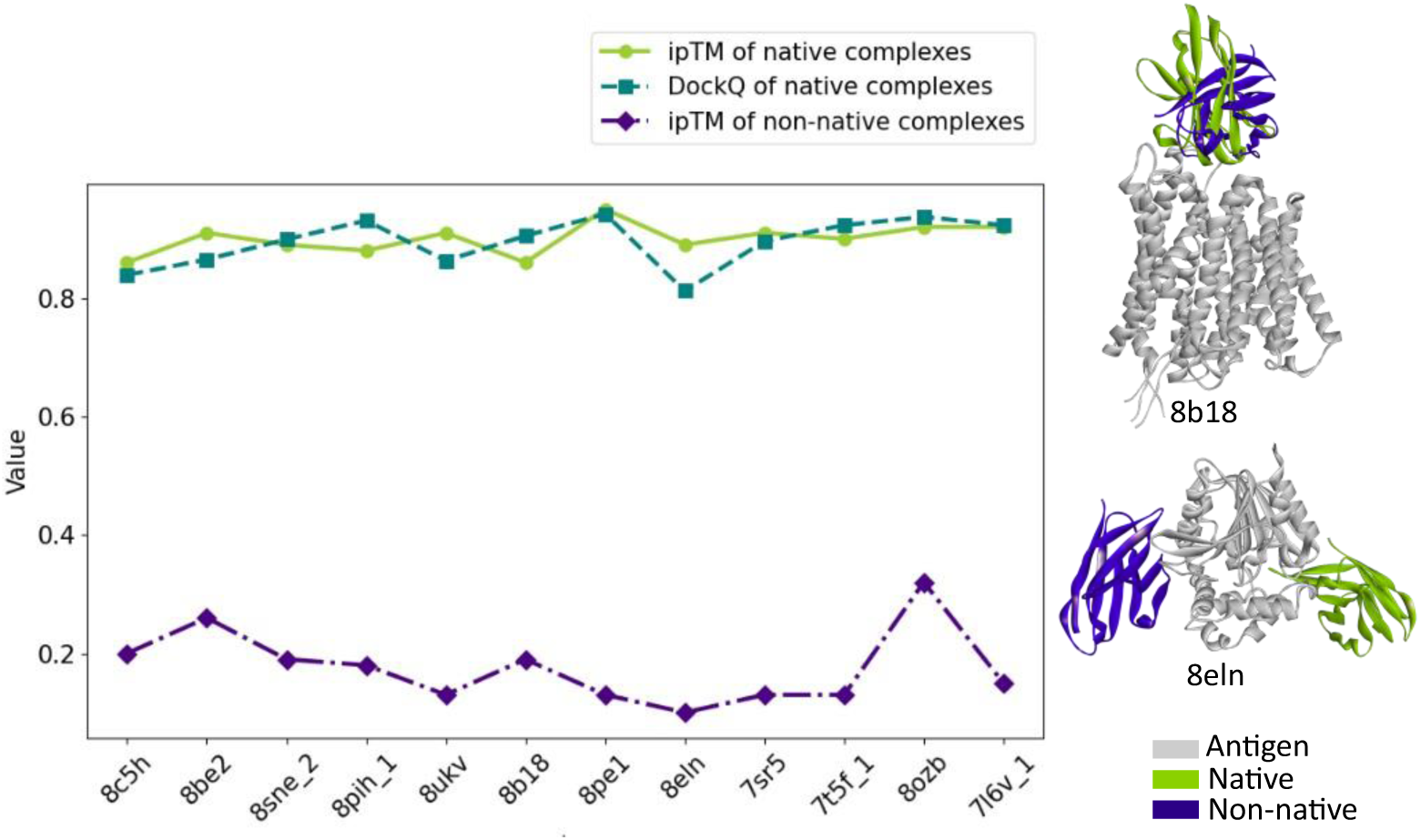
Differentiation between native and non-native complexes using the ipTM score. The x-axis represents the complex examples where the DockQ score was high, indicating a high-quality prediction, and the ipTM score was also high, indicating a confident prediction. The y-axis shows the values for DockQ (orange), ipTM for native complexes (blue) and ipTM for non-native complexes (green). Two illustrative examples of native (green) and non-native (indigo) nanobody epitope prediction against a common antigen (grey) are shown.

Overall, these findings suggest that AF3’s ipTM score is a robust metric for assessing epitope prediction accuracy, with only a few instances in which it did not align with actual prediction performance.

### B. ASSESSING THE ACCURACY OF AF3’S PTM SCORE IN NANOBODY-ANTIGEN COMPLEX PREDICTION

The pTM score which is an estimate of the complex TM-score, ranges from 0 to 1, with scores above 0.5 indicating that the overall predicted fold of the complex and the hypothetical true structure are similar. To evaluate AF3’s accuracy in predicting the pTM score, compared to the true complex TM-score, a strong Pearson correlation coefficient of 0.82 (p-value < 0.001) was calculated, indicating a robust prediction ability of AF3, as represented in Supplementary Information Figure 10.

**Figure 10:**
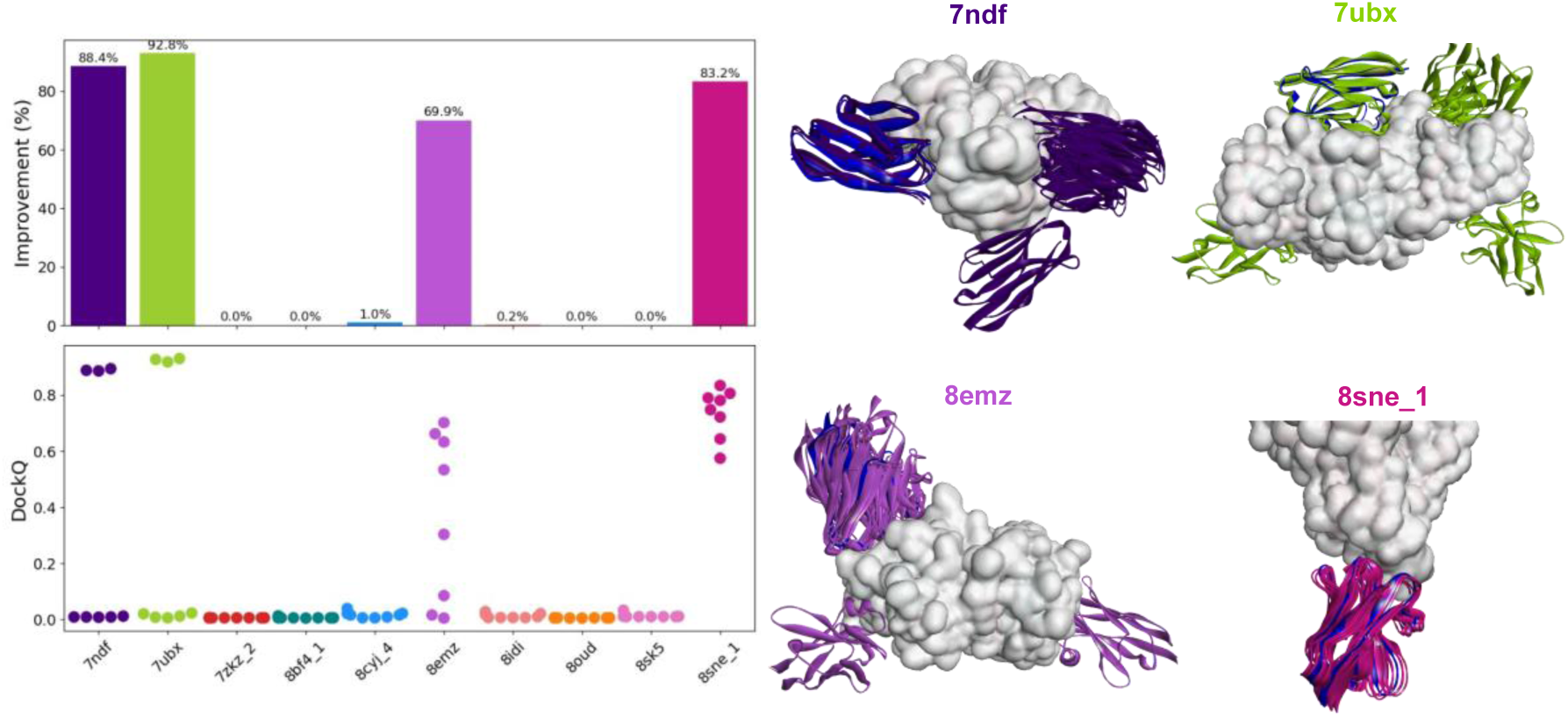
The effect of iterative generation of nanobody/antigen complexes on epitope identification performance. (Left Panel) This dual plot shows PDB examples along the x-axis. The upper plot represents the percentage improvement for each PDB example, based on the DockQ score of the model with the highest ipTM value. The lower plot displays DockQ values on the y-axis across the top ranked model across the 8 runs. (Right Panel) An illustrative example of improved epitope identification, where the crystal structure is colored in blue and the top ranked pose for model seed is displayed.

The strong correlation between the predicted and actual complex TM-scores, as well as the ipTM and DockQ values, further supports the reliability of AF3’s confidence metrics.

### C. UNVEILING AF3’S ABILITY TO IDENTIFY NON-NATIVE NANOBODY-ANTIGEN COMPLEXES

As the accuracy of AF3’s confidence metrics, specifically ipTM, was demonstrated, we further investigated its potential to flag non-native complexes.

High-quality complex predictions with high ipTM scores were selected, and based on the selected complexes, non-native complexes were generated to challenge AF3’s ability to identify them. For each antigen, a non-naturally binding nanobody was paired to create non-native complexes. We found that, AF3, which predicted naturally occurring complexes with high accuracy, assigning them high ipTM values consistent with high DockQ scores, consistently failed to predict non-natural complexes, assigning them low ipTM scores, as illustrated in Figure 9. Interestingly, in some instances, such as 8b18, AF3 was able to identify the epitope despite the presence of a non-native nanobody, while in other cases, such as in 8eln, it identified another epitope. These findings demonstrate that AF3 can effectively discriminate between native and non-native complexes.

In conclusion, AF3’s confidence metrics not only accurately reflect the quality of epitope predictions but also effectively differentiate between native and non-native complexes, opening the door to a broad range of applications.

### V. POTENTIAL OF IMPROVING COMPUTATIONAL EPITOPE IDENTIFICATION

Although AF3 showed enhanced performance over AF2-M in nanobody epitope identification, it remains a significant challenge, with an overall failure exceeding 50%, leaving room for further improvement. We further investigated several alternative and complementary strategies aimed at improving the identification of incorrectly predicted nanobody epitopes.

### A. EVALUATING THE EFFECTIVENESS OF ITERATIVE GENERATION OF NANOBODY-ANTIGEN COMPLEXES ON EPITOPE IDENTIFICATION PERFORMANCE

In this study, we applied an iterative strategy to generate eight model seeds. We found that 4 out of 10 initially inaccurately predicted models showed improvement in epitope prediction, as illustrated in Figure 10. Specifically, 7ndf, 7ubx, 8emz and 8sne_1, exhibited improvements of over 80% for the top ranked pose according to their corresponding ipTM. This improvement may be due to a wider sampling which led to the exploration of the near-native epitopes. However, in most of the cases, 8 generated model seeds were not sufficient for improved predictions. Regarding the accuracy of their ipTM values, we found that the accuracy for classifying correct predictions is 78.75%, as depicted in Supplementary Information Figure 11a.

**Figure 11:**
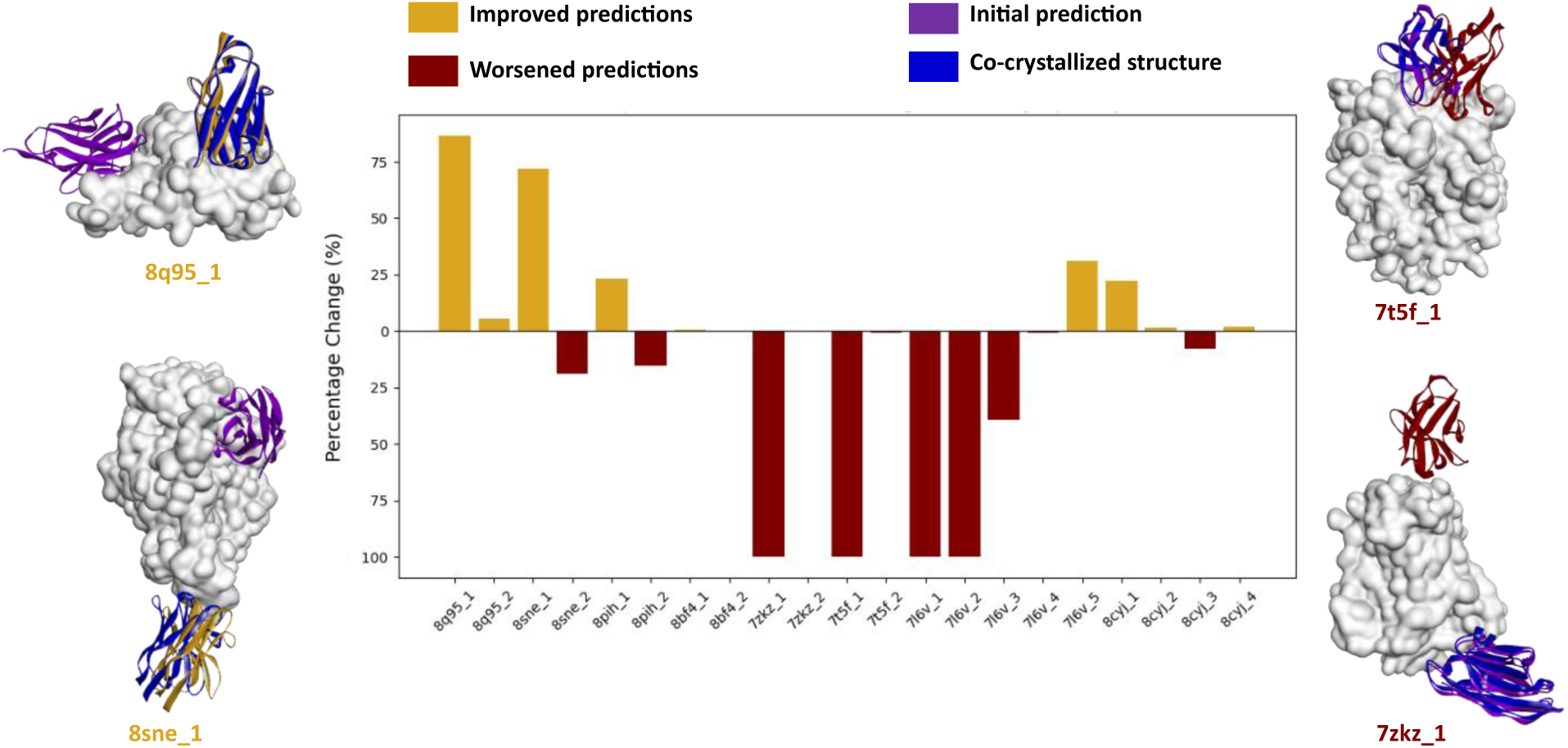
The effect of adding additional co-crystalized nanobodies on the epitope identification performance. This dual plot represents a bar chart showing the percentage change when providing all co-crystalized nanobody sequences for epitope identification using AF3. The y-axis represents the percentage change, with the upper panel representing the percentage improvement (gold) of epitope prediction and the lower panel showing the percentage decline (maroon). The x-axis lists the corresponding multiple epitopes examples. Illustrative examples are provided for improved and worsened predictions.

AF3 was able to generate accurate ipTM with high precision for certain cases, such as 7ubx, but exhibited lower accuracies for others, as shown in Supplementary Information Figure 11b. Additionally various poses densities were observed in case of 8sne_11; all poses were located at the near-native epitope whereas in other examples such as 7ubx, several binding conformations were adopted.

### B. THE INFLUENCE OF ADDITIONAL EPITOPE-RELATED DATA ON EPITOPE IDENTIFICATION PERFORMANCE

In cases where a single antigen is co-crystallized with multiple nanobodies, previous approaches in this study predicted the epitope of each nanobody individually. To improve AF3’s prediction performance, we incorporated all co-crystallized nanobodies to evaluate the impact of additional information on AF3’s prediction capacity. This approach would allow AF3 to explore all potential epitopes and better fit their respective paratopes.

We found that in a few instances, providing additional nanobody sequences resulted in significant improvements in prediction accuracy. However, in most cases, the predictions either remained unchanged or worsened, as illustrated in Figure 11. This may be due to ipTM and pTM values, which reflect the entire antigen complexed with multiple nanobodies, potentially introducing noise and leading to biased rankings.

These findings suggest that while adding additional co-crystallized nanobody information can in some cases improve prediction accuracy, it does not consistently improve epitope identification. In fact, in the majority of cases, it may even reduce the success rate.

## C. THE IMPACT OF RESCORING AF3 MODELS WITHS A PHYSICS-BASED SCORING FUNCTION

In an effort to improve AF3’s nanobody epitope identification, we evaluated the rescoring of the five models generated by AF3 using a robust physics-based scoring function, ZRank, which has demonstrated successful epitope identification in several cases.^44^ However, we found that AF3’s ranking was more accurate than rescoring with ZRank, as shown in Supplementary Information Figure 12.

This shows that even though ZRank is a robust scoring function that showed high accuracy for predicting the near-native epitope among ZDock generated poses, when coupled with AF3 generated poses it was not the optimal approach.^3,4^

An alternative approach for epitope mapping of nanobodies is to use physics-based methods, such as docking predicted structures by homology modeling with tools such as Modeler and ZDock, which have previously been reported for accurate epitope identification of nanobodies.^2–4,45,46^

## MATERIAL AND METHODS

### DATASET SELECTION

### FILTERED SABDAB DATASET

The dataset, downloaded from SabDab as of the July 1, 2024, initially included a total of 1290 bound nanobody structures.^33^ Synthetic constructs, structures with missing residues in their CDRs, and nanobodies exceeding 200 amino acids were removed, resulting in a filtered dataset of 1018 nanobodies. The PDB files were processed using Biopython.^47^ This dataset served as the reference for comparison with the benchmarked dataset.

### BENCHMARKING/SELECTED DATASET

To select the benchmarking dataset, a resolution threshold of 3.6 Å was applied. Structures published recently that were not included in the training datasets for AF3 and AF2-M were chosen. Complexes in which a single nanobody recognized multiple epitopes or had missing residues at the binding site were excluded. However, antigens with several epitopes, co-crystallized with different nanobodies, were assigned the same PDB name with a numerical suffix. A total of 70 nanobody-antigen complexes were manually selected to ensure a diverse representation of antigen types, CDR3 lengths, and CDR3 conformation. Nanobodies were numbered according to Chothia using ANARCI.^48^

### EPITOPE IDENTIFICATION MODELS GENERATION

### ALPHAFOLD2-MULTIMER (VERSION 2.3.2)

AF2-M models were generated using the ‘Predict Protein Structures’ protocol in Discovery Studio Simulation Client.^49^ For MSA generations, the default parameters were set and a total number of recycles of 20 was used. Five generated models on the cloud version of Discovery Studio were ranked according to the AlphaFold ranking system using ipTM and pTM values based on the following formula:

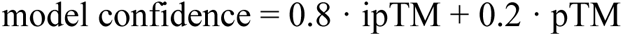

### ALPHAFOLD3

AF3 models were generated using the AlphaFold server. Five generated models were ranked according to the following formula:

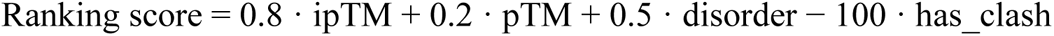

For generating Additional structure models, different run seeds were assigned. For generating the non-native complexes, non-nanobody sequences were added to random receptors present in the dataset.

### EPITOPE IDENTIFICATION EVALUATION AND STRUCTURAL SIMILARITY METRICS DOCKQ.V2

DockQ^39^ assesses the quality of a docked complex with regard to the following equation:

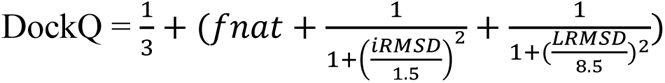

DockQ scores below 0.23 indicate incorrect predictions, scores between 0.23 and 0.49 represent acceptable predictions, scores from 0.49 to 0.80 denote medium quality and scores of 0.80 or above are considered high quality, as represented in Supplementary Information Table 2.

### TM-SCORE

The TM-score ranges from 0 to 1, with a score of 1 indicating a perfect match between both structures. Scores above 0.5 suggest similar folding, while scores above 0.9 indicate near-native structures.^43,50^ TM-score was calculated using tmscoring.

### CDR3 CONFROMATION

CDR3 conformation class was assigned to kinked or stretched based on calculating the pseudo bond and the pseudo dihedral angles (α101 and τ101) of the Cα atoms at the C-terminal end of CDR3.^14,38^ The residues are selected based on the nearly conserved tryptophan located at the C-terminal end of CDR3, assigned number 103, and the three preceding residues: 100, 101 and 102. The classification of CDR3 as kinked or stretched was based on the angles α101 (between residues 100, 101,102 and 103) and τ101 (between residues 100,101 and 102). kinked is when 85°<τ101<130° and 0<α101<120° while stretched is when 100°<τ<120° and -100°>α>101.^14^ However, due to the large conformational space of CDR3 in VHH, exceptions may arise, such as shifts in the kink or the absence of the conserved tryptophan, which can lead to misclassification. Given the size of our dataset, such exceptions were classified based on visual inspection.

### SECONDARY STRUCTURE

The secondary structure was calculated using DSSP, through the Biotite.^51,52^

### DISULFIDE BRIDGES

The disulfide bridge was predicted using the Biotite library by setting a distance of 2.05 Å ±0.07 between sulfur atoms belonging to two cysteine residues and a dihedral angle of 90° ±20.^51^ Values were selected based on the co-crystallized structures.

### STATISTICS

Spearman and Pearson Correlation coefficients were computed using scipy.stats.^53^

### NANOBODY PHYSIOCHEMICAL PROPERTIES

The solvent accessible solvent was calculated using FreeSASA. The additional properties such as Solubility score, Aggregation score, Viscosity score, Developability index and Negative and Positive QMap score were computed using the Calculate Protein Formulation Properties protocol in Discovery Studio 2024. pH was set to 7 and ionic strength to 0.145 using CHARMm forcefield.

### BINDING INTERFACE PROPERTIES

The binding interface properties were computed using the “Analyze Protein Interface” protocol in Discovery Studio 2024.^49^

### ZRANK

ZRANK was computed, as implemented in Discovery Studio 2024.^49^

### CONFUSION MATRIX

Accuracy and Precision were calculated using Scikit-learn library.^54^

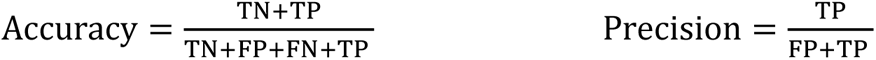

TN: True Negative, TP: True Positive, FN: False Negative, FP: False Positive

## DISCUSSION

As nanobodies emerge as a promising class of biomolecules for therapeutic and diagnostic applications, predicting their epitopes and understanding their binding mechanisms have become a critical step for in silico affinity maturation and epitope-targeted design. The scientific community has long relied on docking experiments to predict nanobody binding modes, a process requiring a high degree of expertise. Recently, epitope identification has become more accessible with the advent of AI-based tools, such as AF2-M and the latest AF3. In this article, we review the performance of both leading algorithms for epitope identification, using a representative and diverse benchmarking dataset of 70 nanobody-antigen complexes.

Despite extensive efforts by the scientific community to enhance the epitope prediction of nanobodies, our findings revealed that both tools successfully identified the binding region with an overall success rate still falling below 50%. This may be attributed to the dynamic nature and selective features of nanobodies, driven by the hypervariability and flexibility of their CDR loops. As CDRs are the key drivers of selective antigen recognition, their flexibility makes predicting a single bound conformation challenging, specifically, with CDR3 reaching considerable lengths and adopting a wide range of conformations.

Despite these findings, AF3 revealed a modest improvement, achieving an overall success rate of 47.1% compared to AF2-M’s 32.8%. However, only a fraction of these predictions qualifies as high-quality models that can be used without further refinement, as they captured nearly all contacts in comparison to the crystal structure. In this context, AF3 nearly doubled the percentage of high-quality predictions, reaching 21.4% compared to AF2-M’s 11.4%. While the remaining predictions successfully identified the binding region, they require further refinement, such as molecular dynamics simulations, to enhance model accuracy and maintain the maximum number of interactions necessary to stabilize the complex.

It has been established that the CDR3 loop is often a key driver of interactions, and its significant variability poses challenges for epitope prediction. Supporting this, both tools performed better when CDR3 exhibited short lengths, as this limits the degree of freedom and consequently reduces its flexibility. Interestingly, our findings indicate that beyond the length of CDR3, its folding orientation whether stretched or kinked, significantly influences the success of epitope prediction. Both tools were more effective when CDR3 adopted a stretched conformation achieving success rates exceeding 50% compared to nearly 20% for kinked. while AF3 struggled to optimize epitope identification for CDR3 adopting kinked conformations, a significant improvement was observed when CDR3 was stretched, with a success rate exceeding 88% compared to 50% for AF2-M. Kinked conformations, which represent the majority of existing nanobodies, continue to pose a significant challenge, affecting the overall performance of AI-driven tools. This may be attributed not only to their unique 3D rearrangement exclusive to nanobodies, but also to how the kinked architecture facilitates the recognition of cryptic epitopes. Additionally, kinked conformations tend to have longer sequences and a higher secondary structure variability compared to stretched conformations.

We also demonstrated that the accuracy of epitope identification strongly relies on the quality of the nanobody model, particularly its CDR3 loop, although high model quality does not always guarantee accurate epitope predictions. Notably, high-quality nanobodies with a stretched conformation significantly increase the likelihood of identifying near-native epitopes. In contrast, the quality of nanobodies adopting kinked conformations is not correlated with the quality of epitope prediction. Furthermore, we found that highly accurate complex models were consistently associated with CDR3 models exhibiting high secondary structure correlation with crystal structures. This indicates that, while RMSD is an important metric, the spatial rearrangement of side chain atoms defined by their secondary structure also plays a critical role in epitope identification.

Regardless of the prediction quality, ipTM proved to be robust and versatile, showing a strong correlation with DockQ results. This suggests that ipTM can serve as a reliable metric for assessing the performance of predicted models. Additionally, we showed that ipTM is sensitive and capable of differentiating between native and non-native complexes, which can be further employed for other applications. Having established the reliability of ipTM, we investigated various strategies to improve epitope identification in poorly predicted complexes. Increasing the run seed and incorporating co-crystallized nanobodies, enhanced the likelihood of identifying near-native epitopes in some instances; however, this enhancement was not consistent, highlighting the complexity of epitope identification and its dependence on numerous factors.

In this study, we have demonstrated that AF3 has made advancements in predicting nanobody-antigen interactions, but particularly excelled in predicting extended conformations, with a success rate exceeding 80%. This represents a remarkable step forward in our ability to model these complex molecular interactions, which are critical for various applications, including therapeutic development.

As the scientific community continues to explore the potential of deep learning in protein interaction prediction, several innovative models are emerging, including Chai-1, HelixFold-Multimer, and OpenFold3, each adopting a unique architecture and offering potential improvements that could further enhance our understanding of protein interactions. Consequently, it would be of great interest to assess these tools regarding their capacity to predict nanobody-antigen complexes predictions while highlighting their respective strengths and limitations.

### ASSOCIATED CONTENT

## Supporting Information

Additional details and data of the benchmarking dataset, the DockQ significance and values across the 70 modeled nanobody-antigen complexes were displayed. In addition, the effect of several factors on epitope identification performance and AlphaFold3’s confidence scores analysis were also presented. Finally, additional strategies for epitope identification improvement were presented.

### AUTHOR INFORMATION

#### Notes

Nanobody® is a registered trademark of Sanofi.

#### DATA AND SOFTWARE AVAILABILITY

PDB files used in this study were downloaded from the Protein databank (https://www.rcsb.org/) and SabDab (https://opig.stats.ox.ac.uk/webapps/SabDab-sabpred/SabDab/nanobodies/) AF3 predictions can be generated using the AlphaFold Server (https://alphafoldserver.com/ ). AF2-M predictions were generated using Discovery Studio Simulation Client.^49^

## ABBREVIATIONS

AF3: AlphaFold3
AF3: AlpaFold-Multimer(v2.3.2), AF2-M
CDR: Complementarity Determining region
FN: False negative
fnat: fraction of correctly predicted reference contacts
FP: False positive
HcAbs: Heavy chain only antibodies
iRMSD: interface root mean square of deviation
ipTM: interface predicted template modeling score
KDE: Kernel Density Estimate
LRMSD: ligand root mean square of deviation
PDB: Protein Data Bank
pTM: predicted Template modeling score
QMap: Surface Charge Map Scores
TN: True negative
TM-score: Template Modeling Score
TP: True positive
VHH: variable heavy domain of heavy chain.

---

Table of Contents Use Only

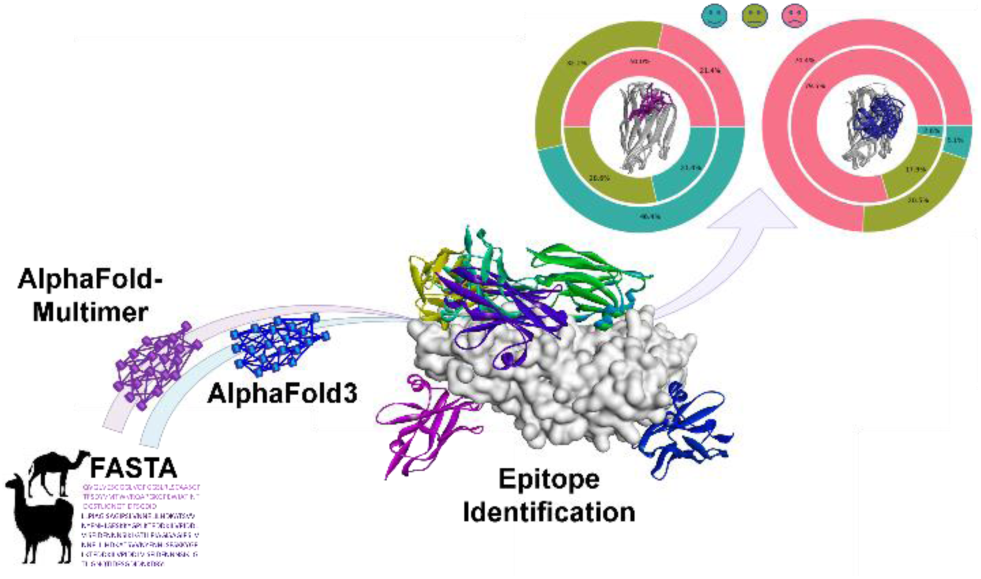

