## Supplementary Information for "AI driven approaches in Nanobody Epitope Prediction: Are We There Yet?"

### Supporting Information Figures:

### SI Figure 1a:

**Diversity within the benchmarking dataset.**

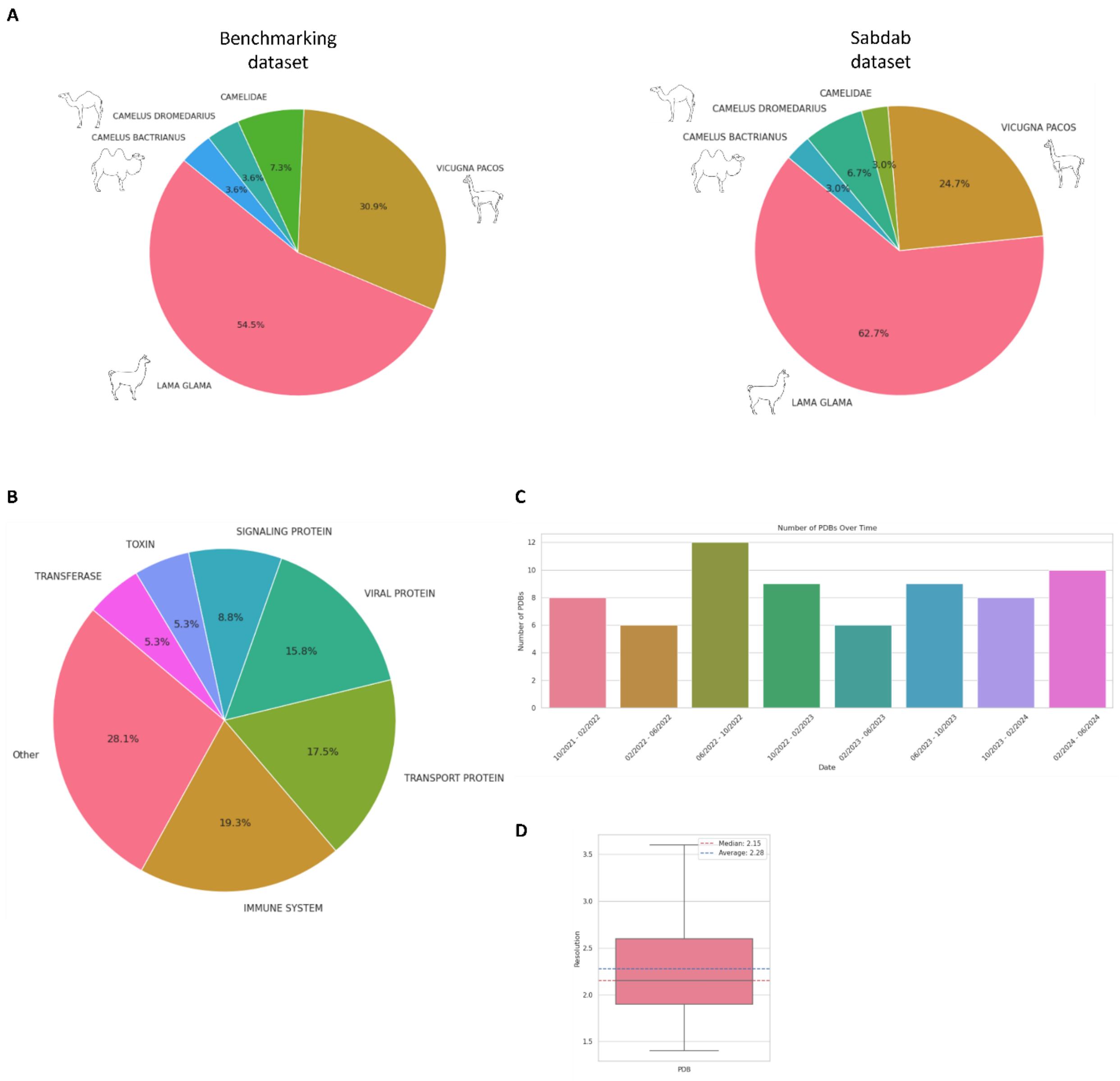

**(A)** <u>The left panel</u> illustrates a pie chart displaying the distribution of nanobody origins represented in the Benchmarking dataset with nanobodies originating from Lama glama accounting for 54.5% of nanobodies. <u>The right panel</u> is a pie chart representing the animal origin of the co-crystallized nanobodies found in the SabDab database. Lama glama represents 62.7% followed by Vicugna pacos at 24.7%. Camelus dromedarius and Camelus bactrianus account for 6.7 % and 3%, respectively. Additionally, 3% of the nanobodies are reported as originating from Camelidae without further specification.

**(B)** A pie chart depicting the protein families to which the antigens belong. Other represents protein families accounting for less than 3%.

**(C)** The distribution of structure release dates for the benchmarking dataset, selected from October 2021 to June 2024, ensuring that the data is recent and was not included in the training datasets of either AlphaFold3 (AF3) nor AlphaFold2-Multimer (AF2-M).

**(D)** The Boxplot represents the resolution distribution of the PDB structures indicating the median and average values.

#### SI Figure 1b:

**Camelidae species from phylogenic tree to CDRs lengths distribution.**

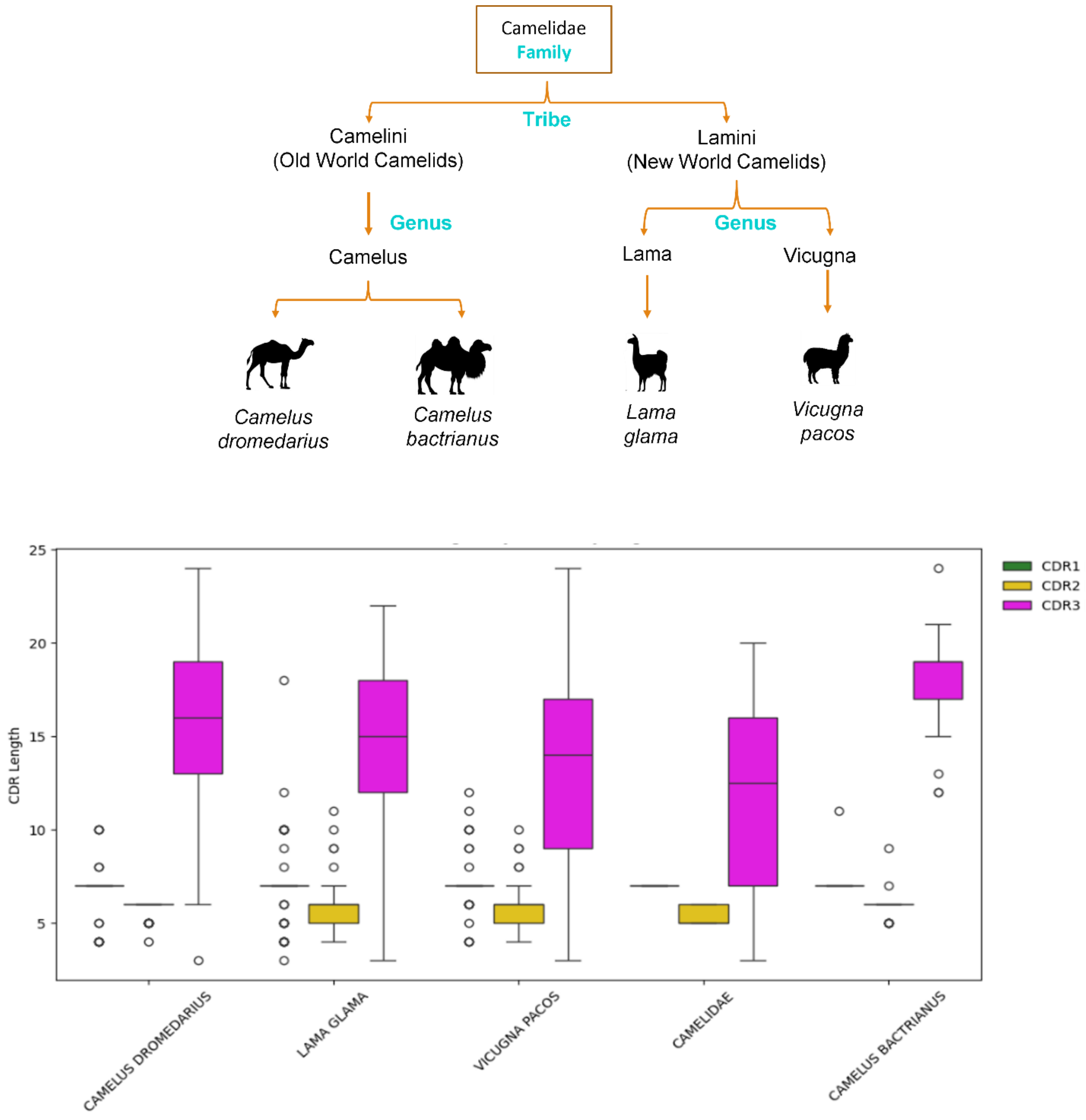

The upper panel is a phylogenic tree of the Camelidae family, illustrating how they evolved into various species with two main tribes: The Old World camelids and New World camelids. Camelus dromedarius and Camelus bactrianus belong to the Old World camelids while Lama glama and Vicugna pacos belong to the New World camelids.

The lower panel is a boxplot representing the distribution of the three CDRs lengths (y-axis) per species (x-axis): Camelus dromedarius, Camelus bactrianus, Camelidae, Lama glama and Vicugna pacos. CDR1 is shown in green, CDR2 in yellow and CDR3 in magenta.

SI Figure 2:

**Interaction differences between CDR3 and corresponding antigen across the different DockQ classes.**

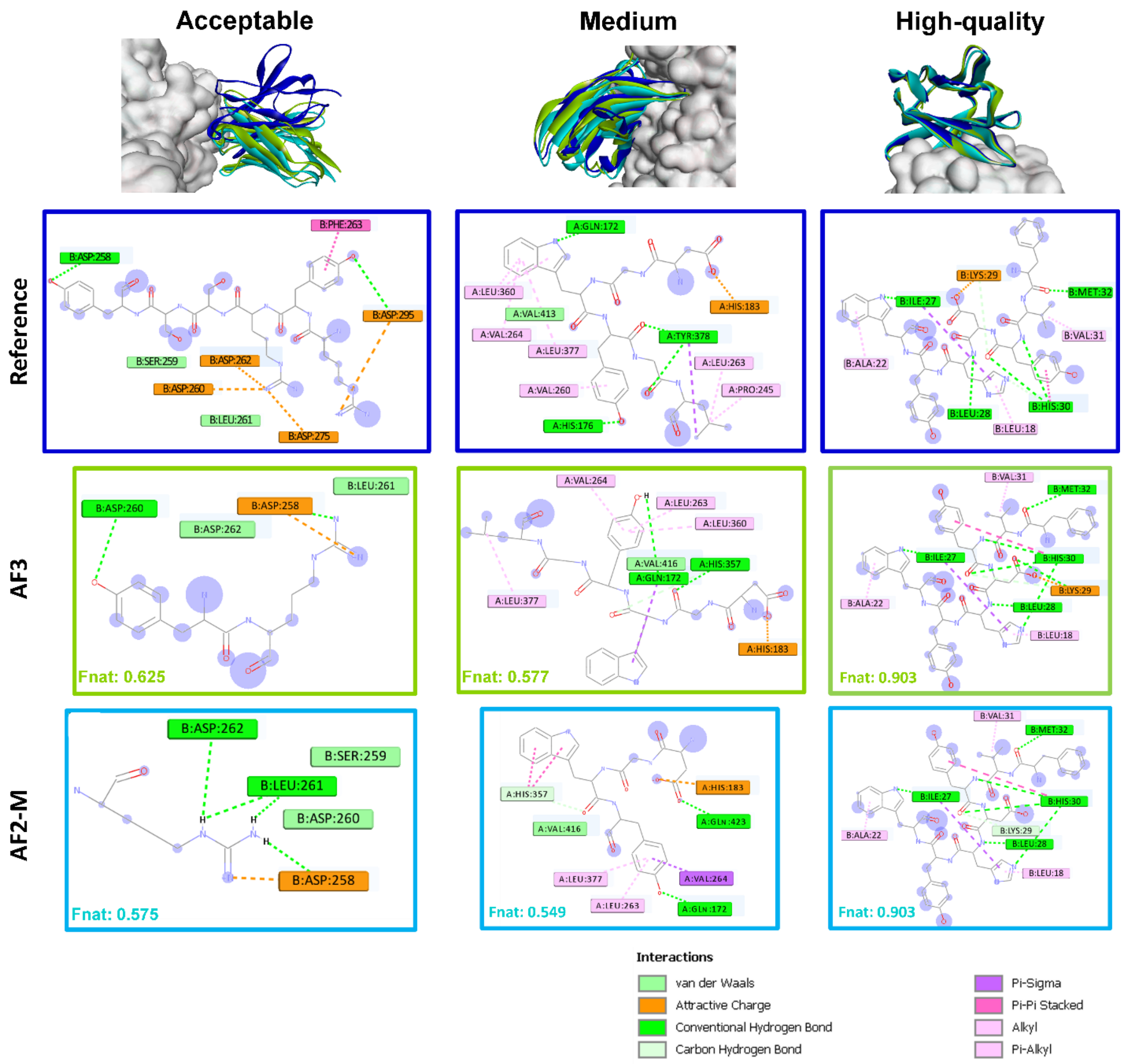

This figure shows the non-bonded interactions (as implemented in Discovery Studio 2024) between CDR3 and the antigen for the AF2-M and AF3 predicted models, compared to the crystal structure across the different DockQ classes.

Protein-protein interactions are crucial when evaluating the epitope identification performances, and in the DockQ score the fraction of native contacts (Fnat) is a value that capture this information. This figure illustrates the degree of retrieved non-bonded interactions between CDR3 and the antigen across various prediction quality levels, ranging from acceptable predictions that preserve a minimal number of interactions to high-quality predictions in which the majority of interactions are maintained across the predicted models.

Acceptable quality predictions: Not all interactions are conserved due to the different orientation of the entire nanobody compared to the reference structure. While the epitope may have been identified, the different backbone orientation of the nanobody makes it interact differently with the antigen, rendering the complex prediction less reliable information and cannot be used as it is. For the example in 7rnn, for both AF3 and AF2-M models, the nanobody was able to interact with Asp258, Asp260, Leu261 and Asp262, similar to the crystal structure, but due to the CDR3 orientation, it was not able to interact with other distant residues such as: Phe263, Asp275, Asp295.

Medium quality predictions: The positioning of the nanobody is more aligned with the crystal structure than in the acceptable quality models; however, the backbone of the CDRs and the side chains are slightly different, preventing the capture of all interactions. In the case of 8et0, both predicted models maintained interactions between CDR3 and the antigen residues: Gln172, His183, Leu377, Leu363, Val264, but failed to maintain interactions with Tyr378 and His176. These missing interactions could potentially be established through molecular dynamics simulations, which might lead the nanobody into a more stable position to retain the most stabilizing contacts in comparison to the crystal structure.

High-quality predictions: They show that both the nanobody backbone and side chains are well oriented, highly matching the crystal structure, preserving the maximum number of interactions between the nanobody and its antigen in both AF3 and AF2-models. Hence this model can be directly used for further calculations.

#### SI Figure 3:

**DockQ values of the 70 predicted complexes by AF3 and AF2-M.**

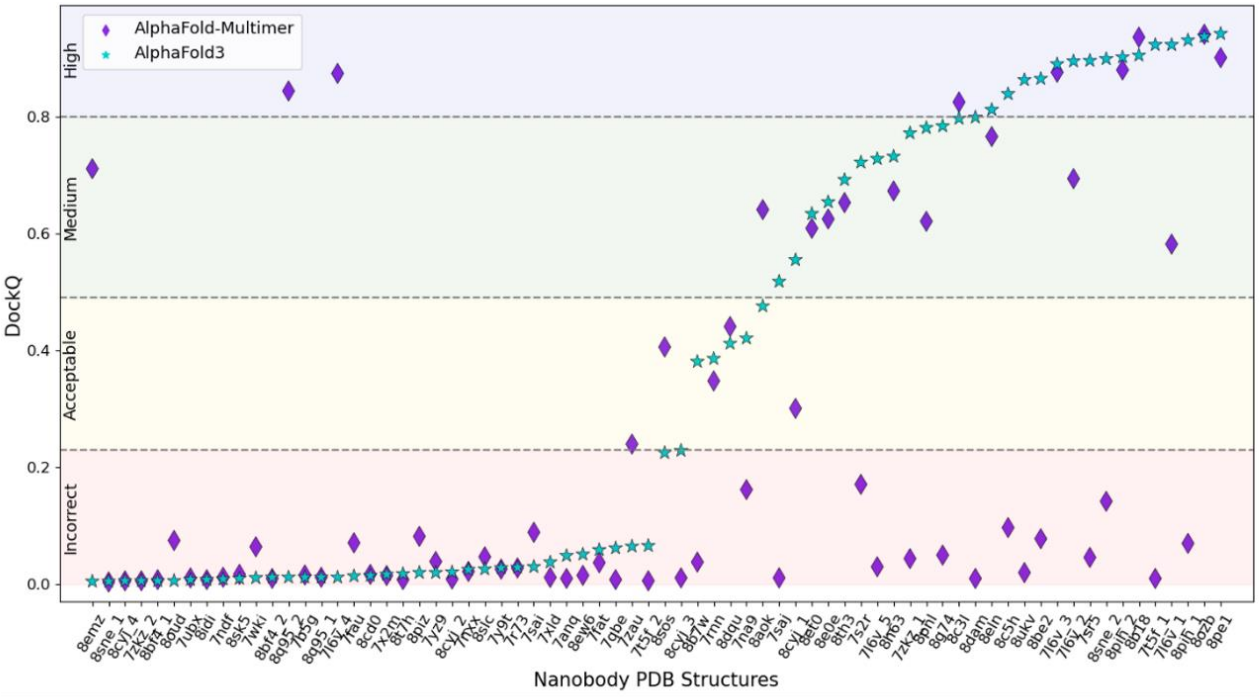

This plot illustrates the performance of both AF3 and AF2-M across 70 complexes. The x-axis represents the 70 different complexes, identified by their PDB ID while the y-axis displays the DockQ scores. The DockQ classification is displayed and is color coded: red for incorrect predictions, yellow for acceptable, green for medium and blue for high quality predictions. AF3 and AF2-M are represented by cyan stars and purple diamonds, respectively. The PDB Ids are arranged in an ascending order of DockQ scores of AF3.

#### SI Figure 4:

##### Effect of CDR3 residue composition on epitope identification performance in AF3.

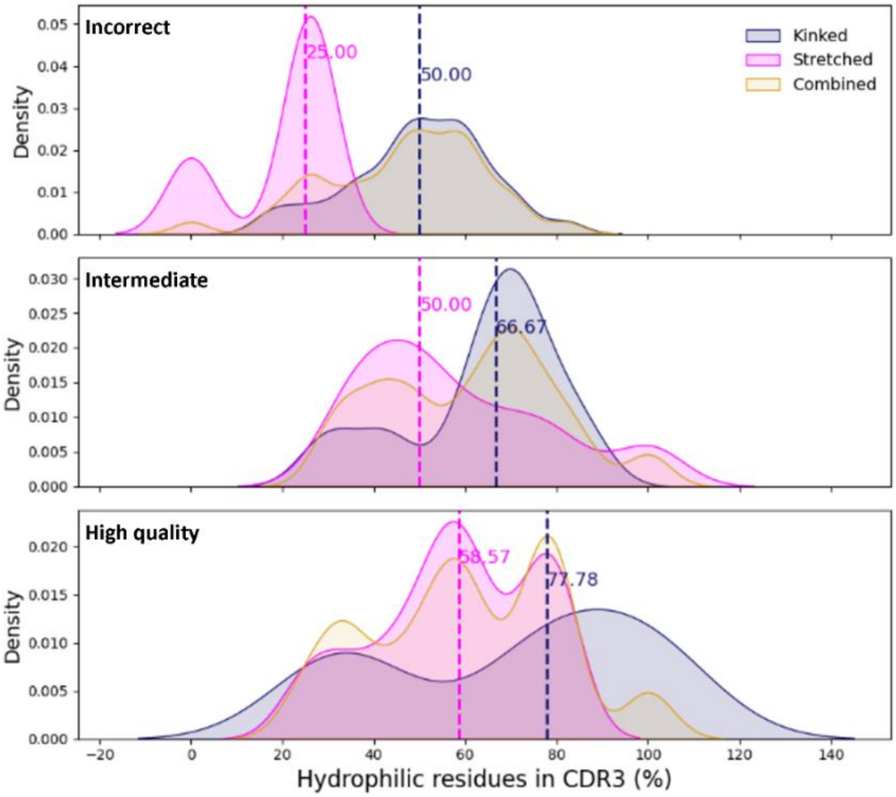

The three plots depict the distribution of hydrophilic residue composition in CDR3 across the three different performance classes for epitope identification: incorrect, intermediate and high-quality. The x-axis represents the percentage of hydrophilic residues among the solvent accessible residues, while the y-axis shows the density distribution. The kinked conformation is represented in dark blue, the stretched conformation in magenta and both conformations combined in gold. Median percentages of hydrophilic residues for each performance class and CDR3 conformation are also displayed.

A shift to the right in the density peak across the DockQ classes is observed, suggesting that a higher proportion of hydrophilic residues in CDR3, as opposed to hydrophobic residues, correlates with better epitope identification performance in AF3. This is further supported by the median percentages of hydrophilic residues in kinked CDR3, which are 50%, 56.67% and 77.78% for the incorrect, intermediate and high-quality DockQ classes, respectively. Similarly, for stretched conformations, incorrect predictions show a median hydrophilic residue percentage of 25%, compared to 58.7% in high-quality poses. In contrast, hydrophobic residues compositions of CDR3 are more frequently associated with lower epitope identification quality. Moreover, a moderate Spearman correlation coefficient of 0.48 was observed between the percentage of hydrophilic solvent accessible residues in stretched CDR3 loops and the DockQ score. However, no significant impact of CDR3 residue composition was observed in AF2-M.

#### SI Figure 5:

**Effect of surface area of the binding interface on epitope prediction performance.**

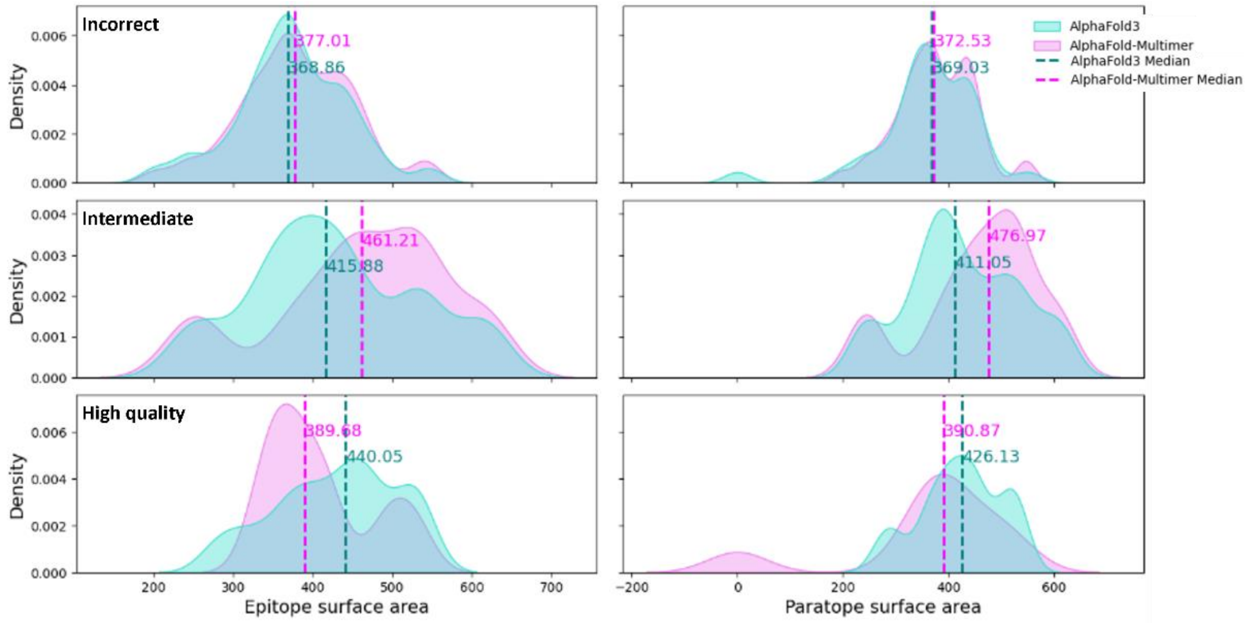

The plots illustrate the distribution of the surface area of the epitope (left panel) and paratope (right panel) across the three performance classes for epitope identification: incorrect, intermediate and high-quality. The x-axis represents the surface area, while the y-axis shows the density distribution. AF3 is colored in turquoise and AF2-M is colored in violet. Median values of surface areas for each performance class are also displayed.

Notably, the median epitope surface area is 368.86 Å for incorrectly predicted epitopes in AF3, compared to 440.05 Å for high quality predictions. Similarly, the median paratope surface area is 369.03 Å in the incorrect predictions, whereas it is 426.13 Å for high-quality predictions. In contrast, AF2-M showed higher median surface areas for the intermediate quality epitope identification but similar median values for both incorrect and high-quality predictions.

#### SI Figure 6a:

**CDR3 conformation classification in AlphaFold3 and AlphaFold-Multimer nanobody (in complex) models.**

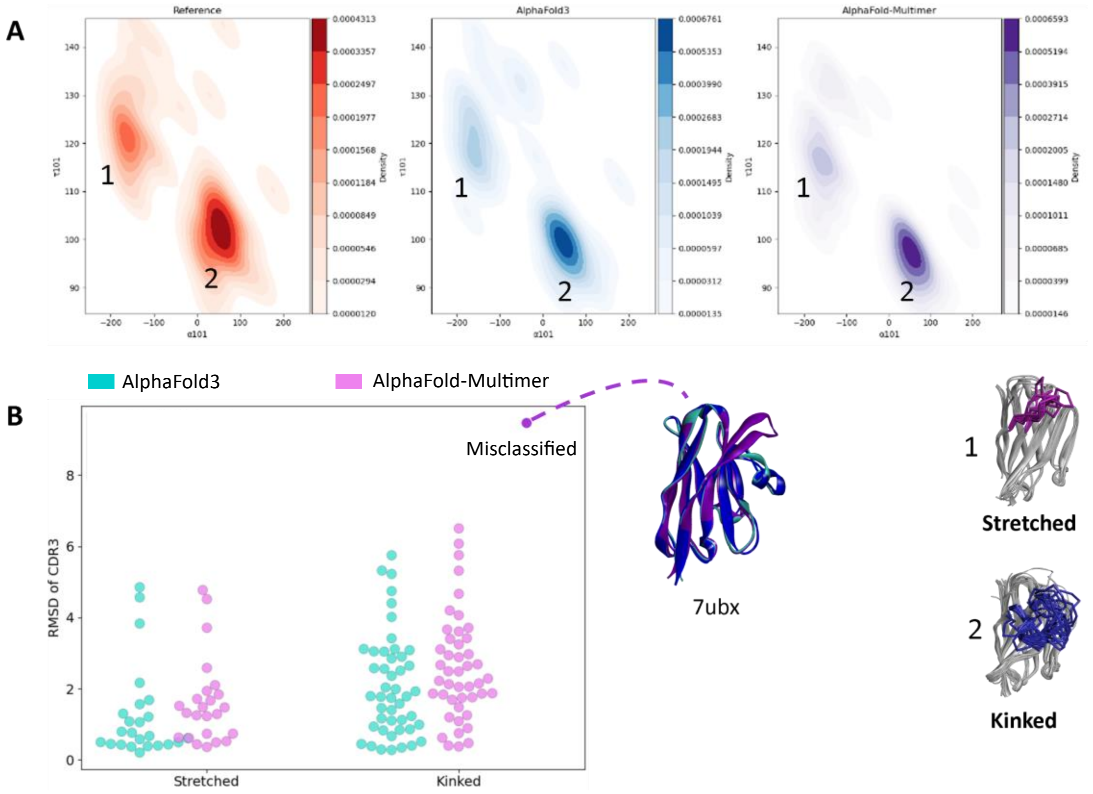

**(A)** This density plot illustrates the distribution of τ101 and α101 angles of CDR3 models in experimental structures (red), AlphaFold3 (blue) and AlphaFold-Multimer (purple). Higher density regions correspond to kinked conformations, while dispersed, less dense clusters represent the stretched conformations. Outliers representing the stretched conformation, reflect the high CDR3 angle flexibility.

**(B)** The swarm plot features the CDR3 conformation (kinked or stretched) on the x-axis and the CDR3 RMSD on the y-axis. AF3 predictions are colored in turquoise whereas AF2-M predictions are colored in violet. Misclassified AF2-M prediction is highlighted and illustrated where the co-crystal nanobody is represented in Blue, AF3 in turquoise and AF2-M in purple.

#### SI Figure 6b:

**Distribution of τ101 and α101 angles of modeled nanobodies versus the co-crystallized structures.**

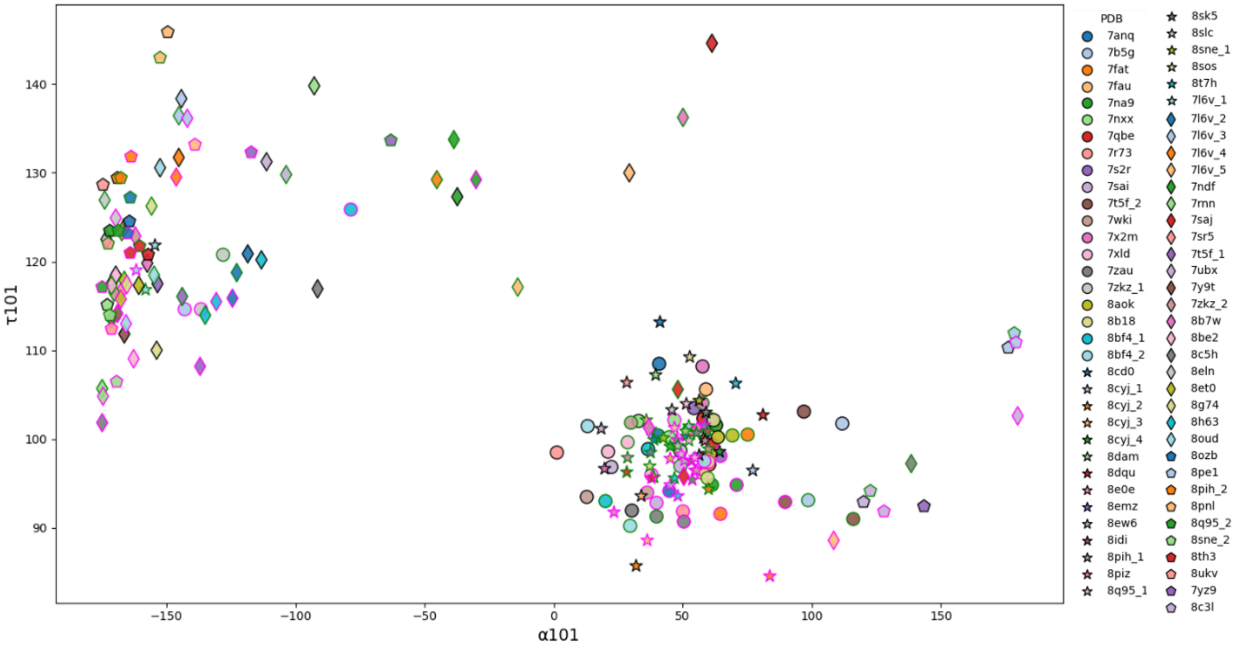

Distribution of τ101 and α101 angles of 70 complexed nanobody in the experimental structure and AlphaFold3 and AlphaFold-Multimer models. Each PDB is given a shape and a color with reference structures are outlined in black, AlphaFold3 is outlined in green and AlphaFold-Multimer is outlined in magenta.

#### SI Figure 7a:

**The secondary structure retrieval percentage of CDR3 models by AlphaFold3 and AlphaFold2-Multimer.**

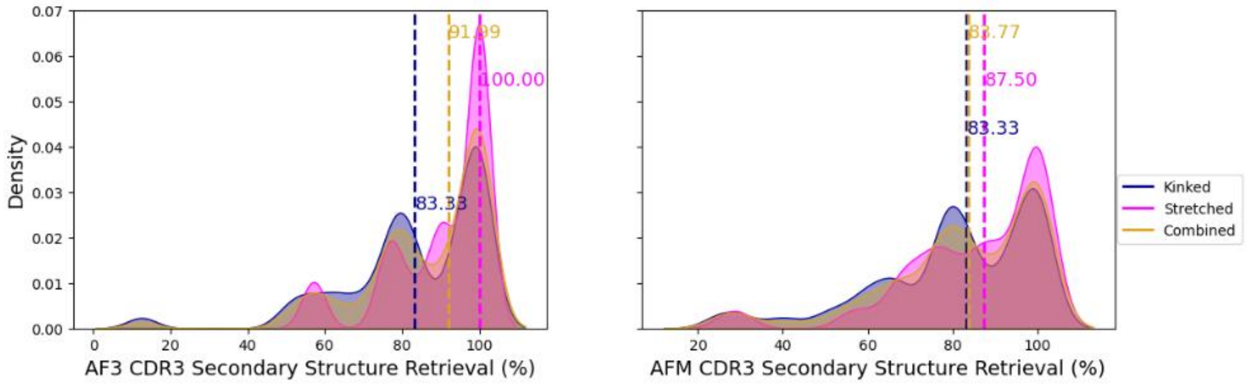

These plots depict the percentage of secondary structure retrieval for the CDR3 regions of modeled nanobodies. The left panel represents the output of AlphaFold3 while the right panel of AlphaFold-Multimer. The kinked conformation is represented in dark blue, the stretched conformation in magenta and both conformations combined in gold. Median percentages for each CDR3 conformation are also displayed.

### SI Figure 7b:

**Effect the secondary structure retrieval of CDR3 on epitope identification performance in AlphaFold3 and AlphaFold2-Multimer.**

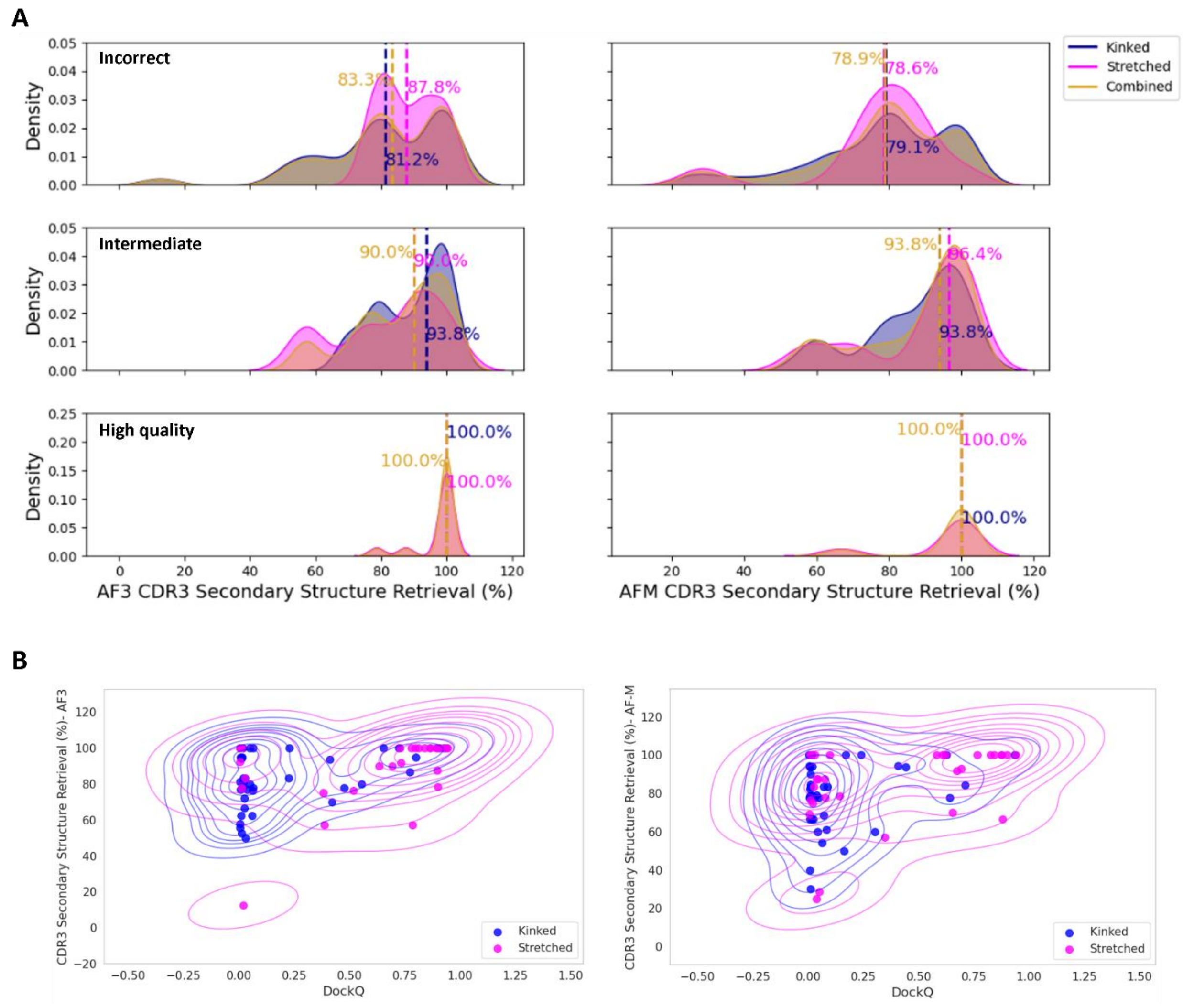

**(A)** These plots depict the percentage of secondary structure retrieval of CDR3 across the three different performance classes for epitope identification: incorrect, intermediate and high-quality. The x-axis represents the percentage of secondary structure retrieval of CDR3, while the y-axis shows the density distribution. The left panel represents the output of AlphaFold3 while the right panel of AlphaFold-Multimer. The kinked conformation is represented in dark blue, the stretched conformation in magenta and both conformations combined in gold. Median percentages of each performance class and CDR3 conformation are also displayed.

**(B)** Scatter plots showing the distribution of conserved secondary structure of CDR3 in percentage on the y-axis versus DockQ score on the x-axis, colored by conformation: stretched (magenta) and kinked (blue). The left panel represents AlphFold3 and while the right panel represents AlphaFold-Multimer. KDE plots are shown to highlight areas of higher density, with denser regions surrounded by more circles. A moderate Spearman correlation between DockQ scores and the retrieval percentage of secondary structure in stretched conformation (0.53 for AF3 and 0.47 for AF2-M with p-values<0.05) was calculated.

#### SI Figure 8:

**Disulfide bridge prediction in nanobody models.**

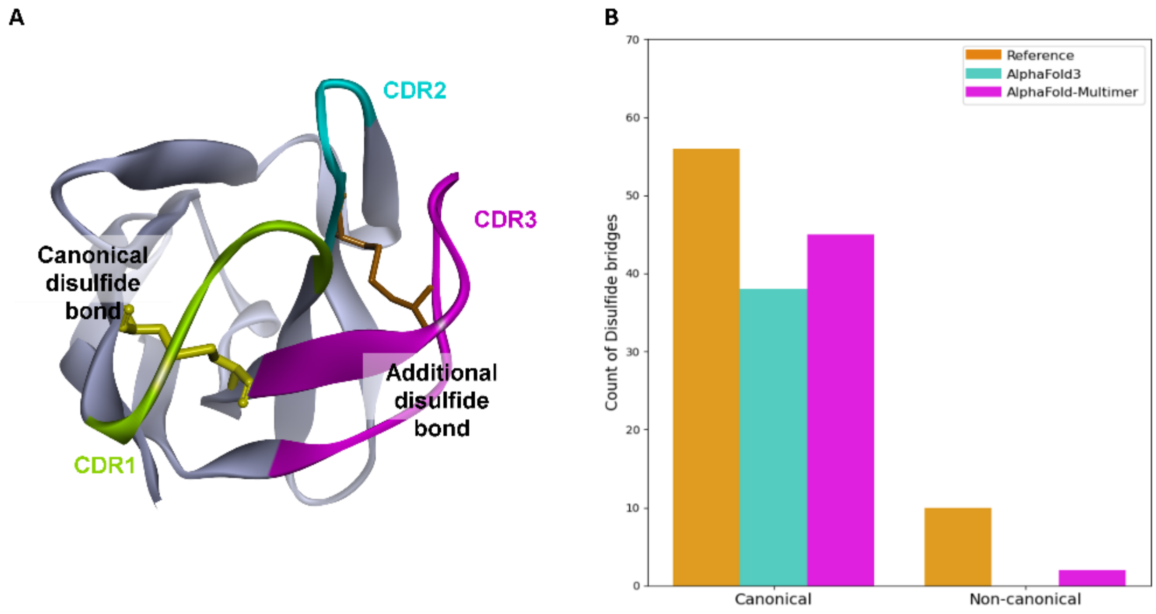

**(A)** Illustration showing canonical and non-canonical disulfide bridges in nanobodies.

**(B)** Bar charts displaying the count of retrieved disulfide bridges on the y-axis versus the type, either canonical or non-canonical on the x-axis. Co-crystallized nanobody structures are represented in orange, AlphaFold3 in cyan and AlphaFold-Multimer in magenta.

#### SI Figure 9:

**Correlation between AF3's confidence score: ipTM and DockQ score.**

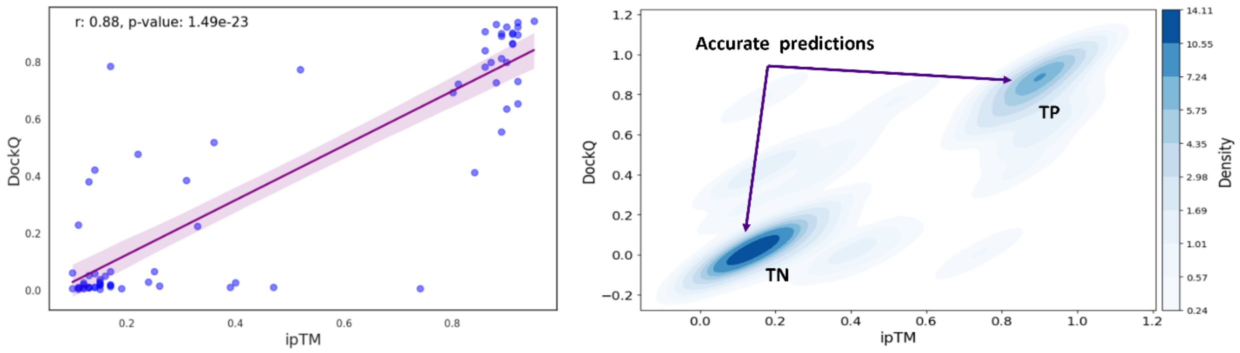

The left panel: it features a scatter plot of the predicted nanobody complex where the x-axis represents ipTM scores and the y-axis represents DockQ scores. A regression line with shaded bands representing the 95% confidence interval, for the regression estimates between the DockQ and ipTM scores is shown. Additionally, Pearson correlation coefficient is calculated along with the p-value.

The right panel: it shows the KDE density distribution, with two denser areas representing cases where both DockQ and ipTM scores align, indicating either high-quality predictions (True Positive, TP) or incorrect ones (True Negative, TN). Misclassified predictions, where ipTM scores do not match the DockQ scores, are represented as less dense clusters.

#### SI Figure 10:

**Spearman Correlation between real complex AF3 TM-score and predicted TM-score (pTM).**

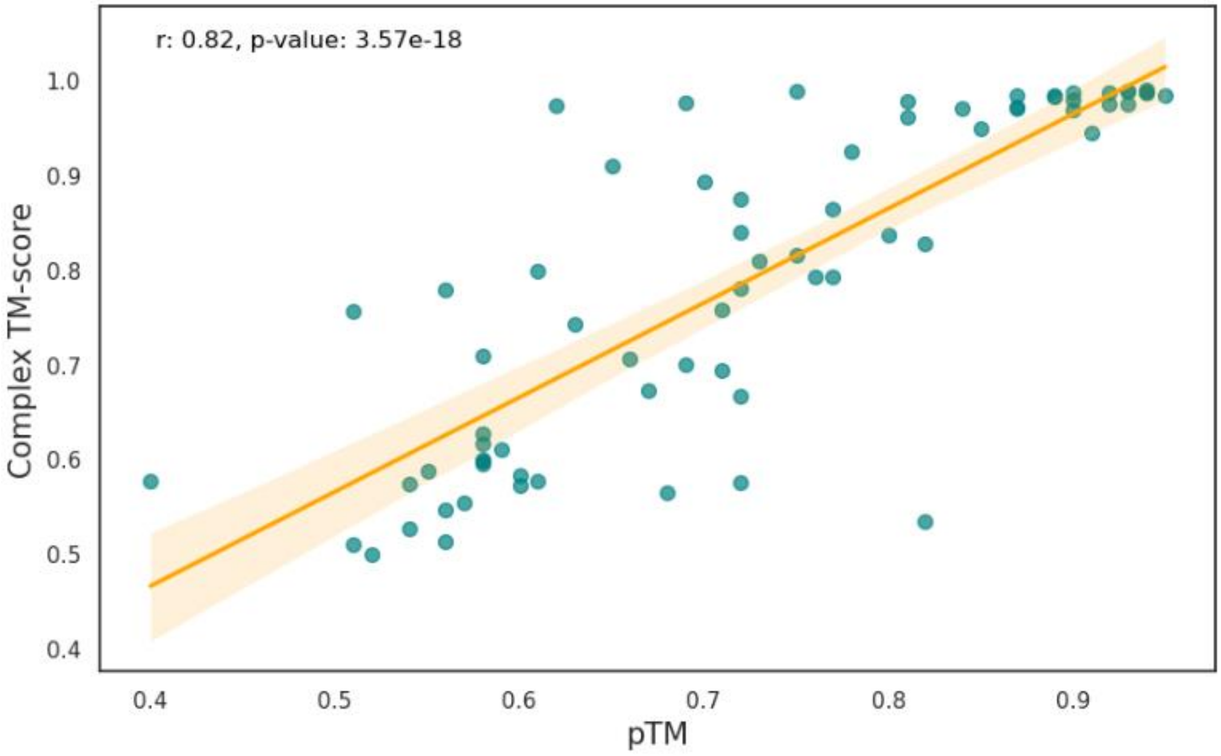

This plot depicts the correlation between the true Complex TM-score on the y-axis and the predicted TM-score on the x-axis. Spearman correlation of 0.73 is shown, denoting a strong correlation between both real and predicted values. A regression line with shaded bands (orange) representing the 95% confidence interval is shown.

#### SI Figure 11a:

**ipTM values across different DockQ classes for iteratively generated models by AF3.**

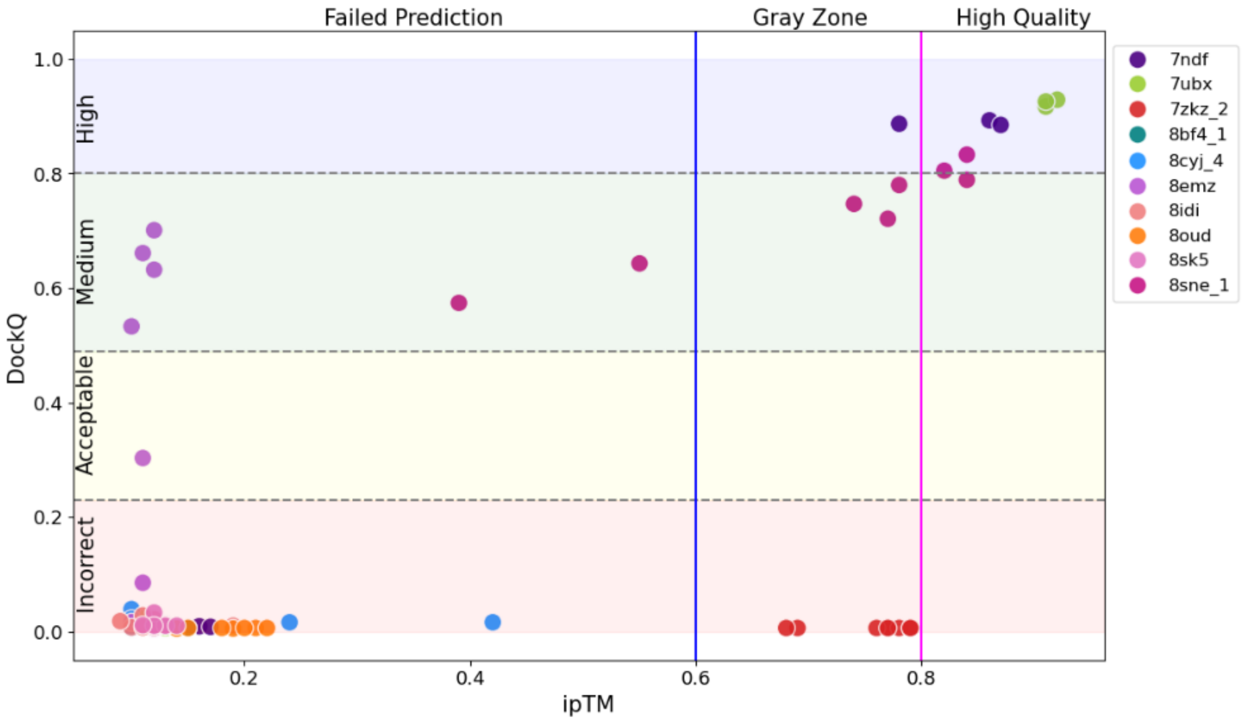

This figure depicts the correlation between the ipTM scores (x-axis) and the DockQ scores (y-axis) of the iteratively generated models by AF3. Each modelled seed is plotted based on its DockQ score and its ipTM score, each complex is represented by a color. DockQ scores are divided into four classes: incorrect, acceptable, medium and high quality, while ipTM scores are classified into three categories: failed prediction, gray zone and high quality.

#### SI Figure 11b:

##### Confusion matrix of ipTM values across DockQ classes for iteratively generated models

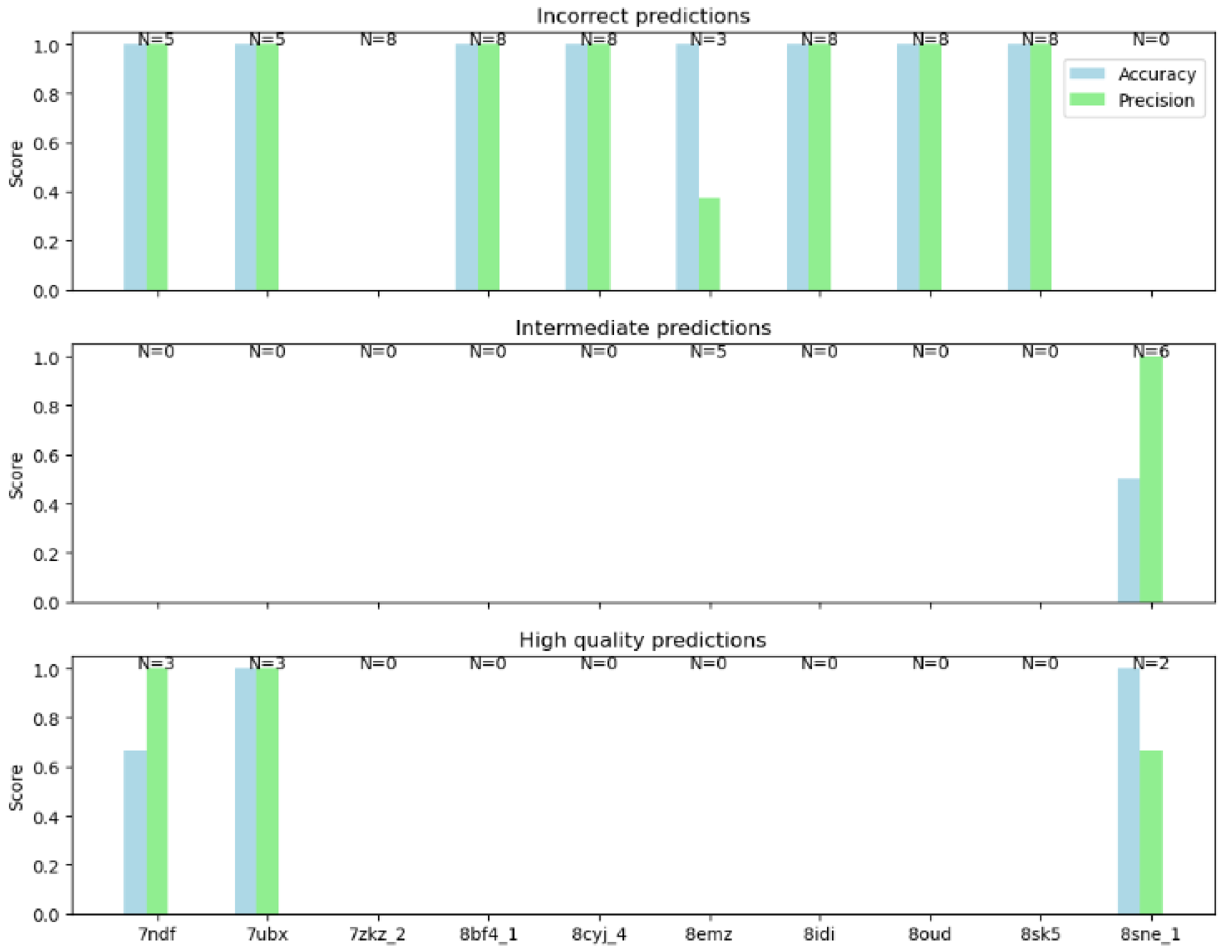

This plot represents a multi-class confusion matrix, where accuracy and precision are displayed as bar charts for each example across different DockQ classes. On the y-axis, score values ranging from 0 to 1 are shown, while on the x-axis are the 10 different examples. Accuracy per example and DockQ class is represented in blue, and precision is represented in green. The number of examples per DockQ class is also indicated.

For instance, in the case of 7ndf, out of the 8 generated model seeds, 5 epitopes were incorrect and 3 were of high-quality. For the 5 incorrect predictions, the ipTM values exhibited high accuracy with high precision. However, for the three high-quality predictions, while precision remained high, accuracy was lower due to the misclassification of one high-quality model.

In contrast, for a case like 7ubx, the ipTM values accurately predicted both incorrect and high-quality models.

In the case of 8emz, 3 were incorrectly predicted with high ipTM accuracy, while 5 were predicted with intermediate quality but were misclassified as failed predictions, resulting in both inaccurate and imprecise ipTM values.

#### SI Figure 12:

##### AlphaFold3 top models selected by two different approaches: ZRank and ipTM

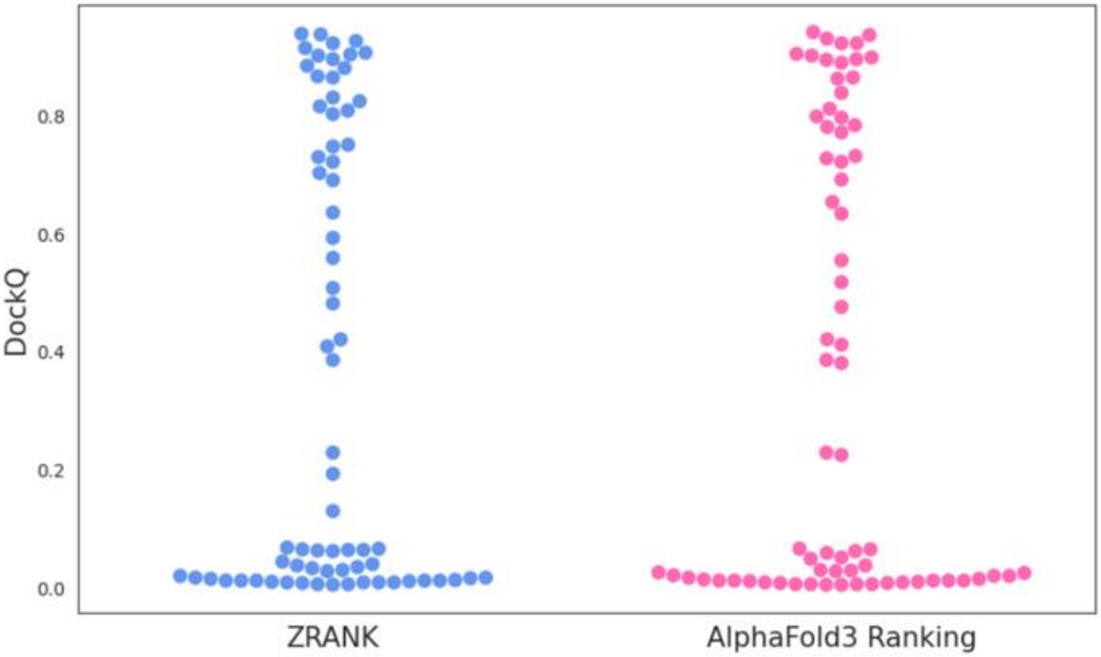

The swarm plot represents DockQ values on the y-axis, comparing the top model according to ZRank rescoring (Blue) and AlphaFold3 ranking (pink). ZRank rescoring takes into account a physics-based approach that accounts for factors such as shape complementarity, hydrophobic contacts, desolvation energy and electrostatic interactions between the nanobody and the antigen. In contrast, AlphaFold3 ranking is guided by the ipTM and pTM scores, which reflect the predicted TM-score for the interface (ipTM) and the entire complex (pTM) between the model and the hypothetical native structure.

### Supporting Information Tables:

#### SI Table 1:

##### Distribution of CDRs lengths of nanobodies through Camelidae species

| CDR1 | MEAN | MEDIAN |
|---|---|---|
| **CAMELIDAE** | 7 | 7 |
| **CAMELUS BACTRIANUS** | 7.15 | 7 |
| **CAMELUS DROMEDARIUS** | 6.86 | 7 |
| **LAMA GLAMA** | 7 | 7 |
| **VICUGNA PACOS** | 7.03 | 7 |
| **CDR2** | | |
| **CAMELIDAE** | 5.46 | 5.0 |
| **CAMELUS BACTRIANUS** | 5.92 | 6.0 |
| **CAMELUS DROMEDARIUS** | 5.84 | 6.0 |
| **LAMA GLAMA** | 5.68 | 6.0 |
| **VICUGNA PACOS** | 5.68 | 6.0 |
| **CDR3** | | |
| **CAMELIDAE** | 12.23 | 13 |
| **CAMELUS BACTRIANUS** | 18 | 19 |
| **CAMELUS DROMEDARIUS** | 16.13 | 16 |
| **LAMA GLAMA** | 14.55 | 15 |
| **VICUGNA PACOS** | 13.37 | 14 |

#### SI Table 2:

**DockQ score thresholds and equivalent model quality**

| **DockQ** | **Model Quality** | |
|---|---|---|
| <0.23 | Incorrect | |
| ≥0.23 | Acceptable | Intermediate |
| ≥0.49 | Medium | Intermediate |
| ≥0.8 | High | |

#### SI Table 3:

##### AF3 and AF2-M predictions of the 70 models across the different DockQ classes

| | Incorrect | Correct | | |
|---|---|---|---|---|
| | | Intermediate Quality | | High Quality |
| | | Acceptable | Medium | |
| **AF3** | **37** | **5** | **13** | **15** |
| **AF2-M** | **47** | **5** | **10** | **8** |

#### SI Table 4:

**Spearman correlation coefficient (ρ) for RMSD of different nanobody regions vs. DockQ scores in epitope identification**

| | Whole Nanobody | All CDRs | CDR1 | CDR2 | CDR3 |
|---|---|---|---|---|---|
| **AF3** | **-0.4526****** | **-0.4796****** | **-0.3481**** | **-0.3212**** | **-0.5264****** |
| **AF2-M** | **-0.4209***** | **-0.4193***** | **-0.1896 (ns)** | **-0.4554****** | **-0.5166****** |

**Weak Correlation** ($0.10< \rho<0.39$) **Moderate Correlation** ($0.40< \rho<0.69$)
ns: no significance *: p-value<0.05 **: p-value<0.01 ***: p-value<0.001 ****: p-value<0.0001

#### SI Table 5:

**Median RMSD values across CDR3 shape classes for both AF2-M and AF3 models**

| | CDR3 (Å) | CDR3 (Å) Kinked | CDR3 (Å) Stretched |
|---|---|---|---|
| **AF3** | **1.34** | **1.781** | **0.761** |
| **AF2-M** | **1.902** | **2.487** | **1.318** |

#### SI Table 6:

**Spearman correlation between the percentage of secondary structure retrieval and the RMSD of CDR3 models in AlphaFold3 and AlphaFold-Multimer.**

| | CDR3 | CDR3 (Kinked) | CDR3 (Stretched) |
|---|---|---|---|
| **AF3** | **-0.618****** | **-0.519***** | **-0.679***** |
| **AF2-M** | **-0.469****** | **-0.464**** | **-0.406*** |

**Weak Correlation** (0.10< ρ<0.39) **Moderate Correlation** (0.40< ρ<0.69)

ns: no significance *: p-value<0.05 **: p-value<0.01 ***: p-value<0.001 ****: p-value<0.0001
